# Parallel HIV-1 fitness landscapes shape viral dynamics in humans and macaques that develop broadly neutralizing antibodies

**DOI:** 10.1101/2024.07.12.603090

**Authors:** Kai S. Shimagaki, Rebecca M. Lynch, John P. Barton

## Abstract

Human immunodeficiency virus (HIV)-1 evolves within individual hosts to escape adaptive immune responses while maintaining its capacity for replication. Coevolution between HIV-1 and the immune system generates extraordinary viral genetic diversity. In some individuals, this process also results in the development of broadly neutralizing antibodies (bnAbs) that can neutralize many viral variants, a key focus of HIV-1 vaccine design. However, a general understanding of the forces that shape virus-immune coevolution within and across hosts remains incomplete. Here we performed a quantitative study of HIV-1 evolution in humans and rhesus macaques, including individuals who developed bnAbs. We observed strong selection early in infection for mutations affecting HIV-1 envelope glycosylation and escape from autologous strain-specific antibodies, followed by weaker selection for bnAb resistance. The inferred fitness effects of HIV-1 mutations in humans and macaques were remarkably similar. Moreover, we observed a striking pattern of rapid HIV-1 fitness gains that precedes the development of bnAbs. Our work highlights strong parallels between infection in rhesus macaques and humans, and it reveals a quantitative evolutionary signature of bnAb development.

## Introduction

Human immunodeficiency virus (HIV-1) rapidly mutates and proliferates in infected individuals. The immune system is a major driver of HIV-1 evolution, as the virus accumulates mutations to escape from host T cells and antibodies ^1–3^. Due to the chronic nature of HIV-1 infection, coupled with high rates of mutation and replication, HIV-1 genetic diversity within and between infected individuals is incredibly high. Genetic diversity challenges vaccine development, as vaccine-elicited antibodies must be able to neutralize many strains of the virus to protect against infection ^4^.

However, there exist rare antibodies that are capable of neutralizing a broad range of HIV-1 viruses. These broadly neutralizing antibodies (bnAbs) have therefore been the subject of intense research ^5–8^. Eliciting bnAbs through vaccination remains a major goal of HIV-1 vaccine design. However, the development of exceptionally broad antibody responses is rare, and such antibodies typically develop only after several years of infection ^9–12^.

Recent years have yielded important insights into the coevolutionary process between HIV-1 and antibodies that sometimes leads to the development of bnAbs. Clinical studies have collected serial samples of HIV-1 sequences from a few individuals who developed bnAbs and characterized the resulting antibodies, their developmental stages, and binding sites ^13–15^. The contributions of HIV-1 and its coevolution to bnAb development are complex ^16,17^. High viral loads and viral diversity have been positively associated with bnAb development ^16–18^. However, superinfection, which can vastly increase HIV-1 diversity, is not always associated with bnAb development ^19^, and it does not appear to broaden antibody responses in the absence of other factors ^17^.

Here, we sought to characterize the evolutionary dynamics of HIV-1 that accompany the development of bnAbs in clinical data. In particular, we inferred the landscape of selective pressures that shape the evolution of HIV-1 within hosts, reflecting the effects of the immune environment. We first analyzed data from two individuals who developed bnAbs within a few years after HIV-1 infection ^13,14^. In both individuals, HIV-1 mutations inferred to be the most beneficial were observed early in infection. In general, mutations that provided resistance to autologous strain-specific antibodies were inferred to be more strongly selected than ones that escaped from bnAbs. We also observed clusters of beneficial mutations along the HIV-1 genome, which were associated with envelope protein (Env) structure.

To confirm the generality of these patterns in a broader sample, we studied recent data from rhesus macaques (RMs) infected with simian-human immunodeficiency viruses (SHIV) that incorporated HIV-1 Env proteins derived from the two individuals above ^20^. This study also compared patterns of Env evolution in HIV-1 and SHIV in response to host immunity. We observed striking parallels between the inferred fitness effects of Env mutations in RMs and humans, suggesting highly similar selective pressures on the virus despite different host species and differences in individual immune responses. Furthermore, we found that RMs that developed broad, potent antibody responses could clearly be distinguished from those with narrowly focused responses using the evolutionary dynamics of the virus. Specifically, the virus population in individuals who developed greater breadth was distinguished by larger and more rapid gains in fitness than in other individuals. Collectively, these results show high similarity between SHIV evolutionary dynamics in RMs and HIV-1 in humans, and that viral fitness gain is associated with antibody breadth.

## Results

### Quantifying HIV-1 evolutionary dynamics

We studied HIV-1 evolution accompanying the development of bnAbs in two donors, CH505 and CH848, enrolled in the Center for HIV/AIDS Vaccine Immunology 001 acute infection cohort ^21^. CH505 developed the CD4 binding site-targeting bnAb CH103, which was first detectable 14 weeks after HIV-1 infection ^13^. CH103 maturation was found to be associated with viral escape from another antibody lineage, CH235, that ultimately developed significant breadth ^22,23^. CH848 developed a bnAb, DH270, targeting a glycosylated site near the third variable loop (V3) of Env ^14^. Similar to the bnAb development process in CH505, escape from “cooperating” DH272 and DH475 lineage antibodies was observed to contribute to the maturation of DH270 (ref. ^14^).

To quantify HIV-1 evolutionary dynamics, we sought to infer a fitness model that best explained the changes in the genetic composition of the viral population observed in each individual over time. In recent years, a wide variety of approaches have been developed to infer the fitness effects of mutations from temporal genetic data ^24–43^. The vast majority of these methods focus on a single locus at a time, ignoring correlations between genetic variants at different loci. While the recombination rate of HIV-1 is high ^44,45^, the virus also evolves under strong selection, which can lead to interference between clones with different beneficial mutations ^39,46–50^. Thus, we applied MPL, an inference method that systematically accounts for genetic correlations ^39–43^, to estimate the fitness effects of HIV-1 mutations.

### Model overview

Here, we provide a brief overview of the key steps in the MPL approach to inferring selection. Further details are available in Methods and in prior work ^39–43^. First, we assume that the effect on viral fitness of each individual mutation *a* at each site *i* is quantified by a selection coefficient *s*_*i*_(*a*), with positive coefficients *s*_*i*_(*a*) *>* 0 denoting mutations that are beneficial for the virus and *s*_*i*_(*a*) *<* 0 denoting deleterious ones. We further assume that the cumulative fitness effects of mutations are additive, such that the overall fitness *F*^*α*^ of a viral sequence *α* is given by the sum of the selection coefficients for all the mutations that it bears. That is,

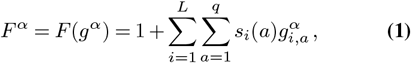

where 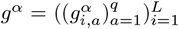 represents the viral sequence, with 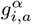 qual to one if genotype *α* has allele *a* at site *I* and zero otherwise. *L* is the length of the genetic sequence, and *q* represents the number of genetic states (i.e., *q* = 5 for nucleotide sequences and 21 for amino acids, including gaps/deletions).

We assume that viral replication is stochastic, where viruses with higher fitness are more likely to spread infection to new cells than ones with lower fitness. Let us write the number of viruses of each genotype in the population at time *t* as *n*(*t*) = (*n*_1_(*t*), *n*_2_(*t*), …, *n*_*M*_ (*t*)), where 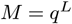 is the total number of possible genotypes. In our model, the probability of obtaining a new distribution of genotypes *n*(*t* + 1) in the next generation is multinomial,

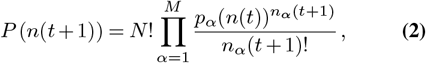

with *N* = ∑_*α*_*n*_*α*_ the total population size. The probabilities *p*_*α*_(*n*(*t*)) are influenced by fitness, as well as mutation, recombination, and the current frequency of each genotype (see Methods). Essentially, viruses that are fitter than the current population average are more likely to increase in frequency, while ones that are less fit are likely to decline. Mutations and recombination introduce genetic variation into the population. The population size *N* determines the relative stochasticity of the dynamics, with smaller populations having more random fluctuations than larger ones.

Following this model, we can then quantify the probability of any evolutionary history of the viral population (i.e., the distribution of viral genotypes over time) as a function of the selection coefficients (Methods). Assuming that the population size *N* is large and the selection coefficients are small (such that |*s*| *≪* 1), we can write an analytical expression for the selection coefficients that best fit the viral dynamics observed in data ^39,40^:

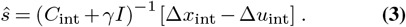

Here *C*_int_ is the covariance matrix of allele frequencies (i.e., the linkage disequilibrium matrix) integrated over time, and *γ* is a regularization parameter. The terms Δ*x*_int_ and Δ*u*_int_ represent the net change in allele frequency and the total expected change in allele frequency due to spontaneous mutations alone, respectively (see Methods for details). Intuitively, this expression says that net allele frequency changes that cannot be explained by mutations are likely due to selection, either on that specific allele or the associated genetic background, which is quantified by *C*_int_. Alleles that have large, rapid changes in frequency are more likely to be under strong selection than those with smaller, slower frequency changes. Beyond HIV-1 (ref. ^39^), this approach has been successfully applied to study the evolution of SARS-CoV-2 (ref. ^43^) and experimental evolution in bacteria ^51,52^.

### Broad patterns of HIV-1 selection

As described above, we used MPL ^39,40^ to infer the selection coefficients that best fit the viral dynamics observed in data from CH505 and CH848. While the great majority of HIV-1 mutations were inferred to be neutral (*s*_*i*_(*a*)*∼* 0), a few mutations substantially increase viral fitness (**Fig. 1**). Strongly beneficial mutations occurred in clusters along the genome and preferentially appeared in specific regions of Env (**Fig. 2**).

**Fig. 1.**
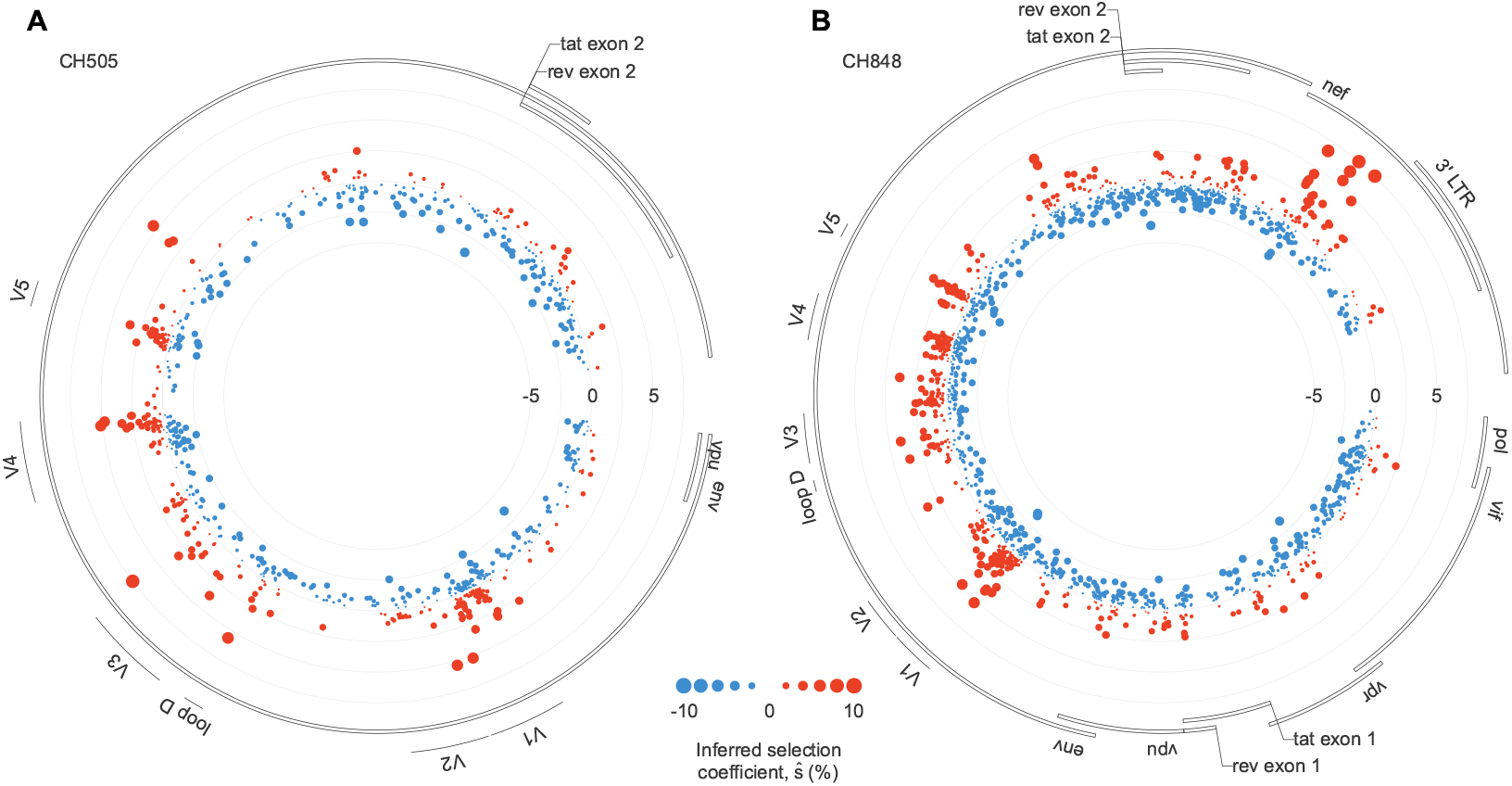
Beneficial mutations occur in clusters along the genome. Inferred fitness effects of HIV-1 mutations in CH505 (**A**) and CH848 (**B**). The position along the radius of each circle specifies the strength of selection: mutations plotted closer to the center are more deleterious, while those closer to the edge are more beneficial. For both individuals, clusters of beneficial mutations are observed in the variable loops of Env, some of which are associated with antibody escape. For CH848, a group of strongly beneficial mutations also appears in Nef.

**Fig. 2.**
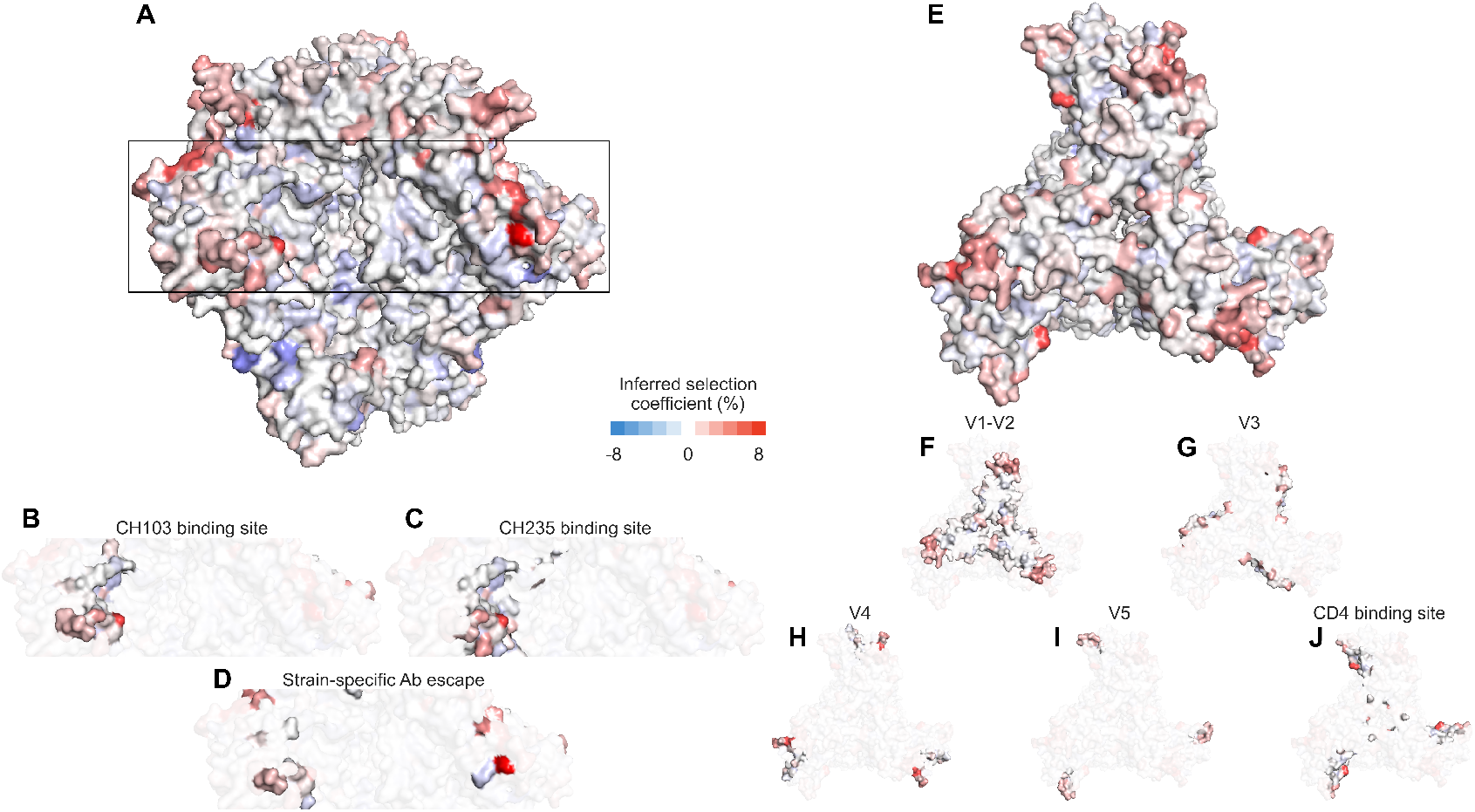
Visualization of the effects of HIV-1 mutations in CH505 on the Env trimer. **A**, Side view of the Env trimer ^53^, with detail views of selection for mutations in the CH103 binding site (**B**), CH235 binding site (**C**), and sites associated with escape from autologous strain-specific antibodies (**D**). **E**, Top view of the trimer, with detail views of variable loops (**F-I**) and the CD4 binding site (**J**). Generally, beneficial mutations appear more frequently near exposed regions at the top of the Env trimer, and deleterious mutations appear in more protected regions. Mutations near the V1/V2 apex can affect the binding and neutralization of antibodies targeting the CD4 binding site ^22^.

To quantify patterns of selection in Env, we examined the top 2% of mutations inferred to be the most beneficial in CH505 and CH848. The fractions of nonsynonymous mutations within these subsets were 97% and 92% for CH505 and CH848, respectively. These fractions are significantly higher than chance expectations (*p* = 6.7 *×* 10^−3^ and 5.6 *×* 10^−4^, Fisher’s exact test; Methods), supporting the model’s ability to accurately infer fitness effects in this data. For CH505, we found 10.9-fold more strongly beneficial mutations in the first variable loop (V1) than expected by chance (*p* = 2.5 *×* 10^−3^). This is consistent with the presence of V1 mutations conferring resistance to autologous strain-specific antibodies ^22^. Mutations in V4, a region targeted by CD8^+^ T cells ^22^, were also 10.0-fold enriched in this subset (*p* = 9.0 *×* 10^−5^). For CH848, mutations in V1, V3, and V5 were enriched by factors of 14.2, 6.3, and 19.1 among the top 2% most beneficial mutations (*p* = 5.9 *×* 10^−10^, 6.6 *×* 10^−3^, and 4.8 *×* 10^−6^). Mutations in these regions were shown to play a role in resistance to DH270 and DH475 lineage antibodies ^14^. To test whether our results might be biased by overall sequence variability, we examined the relationship between our inferred selection coefficients and entropy, a common measure of sequence variability. Overall, we found only a modest correlation between selection and entropy, suggesting that the signs of selection that we observe are not due to increased sequence variability alone (**Supplementary Fig. 1**).

Reversions and mutations affecting N-linked glycosylation motifs were also likely to be beneficial. We define reversions as mutations where the transmitted/founder (TF) amino acid changes to match the subtype consensus sequence at the same site. Among the top 2% most beneficial mutations, reversions were enriched by factors of 19.9 and 17.8 for CH505 and CH848 viruses, respectively (*p* = 2.1 *×* 10^−8^ and 8.5 *×* 10^−13^), consistent with past work finding strong selection for reversions ^39,54^. For CH848, this group also includes several strongly selected mutations observed in Nef (**Fig. 1B**), a protein that plays multiple roles during HIV-1 infection ^55,56^. Mutations affecting N-linked glycosylation motifs (i.e., by adding, removing, or shifting a glycosylation motif) were enriched by factors of 4.6 (*p* = 7.0 *×* 10^−3^) and 8.7 (*p* = 1.4 *×* 10^−10^). Changes in glycosylation patterns contributed to antibody escape for both CH505 and CH848 (refs. ^14,22^).

### Selection for antibody escape

To quantify levels of selection for antibody escape, we computed selection coefficients for mutations that were observed to contribute to resistance to bnAbs as well as bnAb precursors and autologous strain-specific antibodies (**Supplementary Tables 1-3**) ^22,57–60^. We mapped the inferred selection coefficients to the Env protein structure ^13^, highlighting the binding sites for bnAbs and resistance mutations for strain-specific antibodies, as well as important parts of Env (**Fig. 2, Supplementary Fig. 2**). We also analyzed when these resistance mutations were first observed in each individual.

Overall, we observed stronger selection for escape from autologous strain-specific antibodies and/or changes in glycosylation during the first six months of infection (**Supplementary Tables 4-5**). This was then followed by more modest selection for escape from bnAb lineages (**Fig. 3**). For CH505, mutations that conferred resistance to the intermediate-breadth CH235 lineage ^22^ were less beneficial than top mutations escaping from strain-specific antibodies (**Fig. 3A**). In turn, resistance mutations for the broader CH103 lineage were less beneficial than CH235 resistance mutations (**Fig. 3A**). CH848 is similar, with some highly beneficial mutations affecting glycosylation observed early in infection (**Fig. 3B**). While a few mutations affecting DH270 appear strongly selected, these mutations appeared long before the DH270 lineage was detected (around 3.5 years after infection ^14^). Thus, these mutations may have initially been selected for other reasons.

**Fig. 3.**
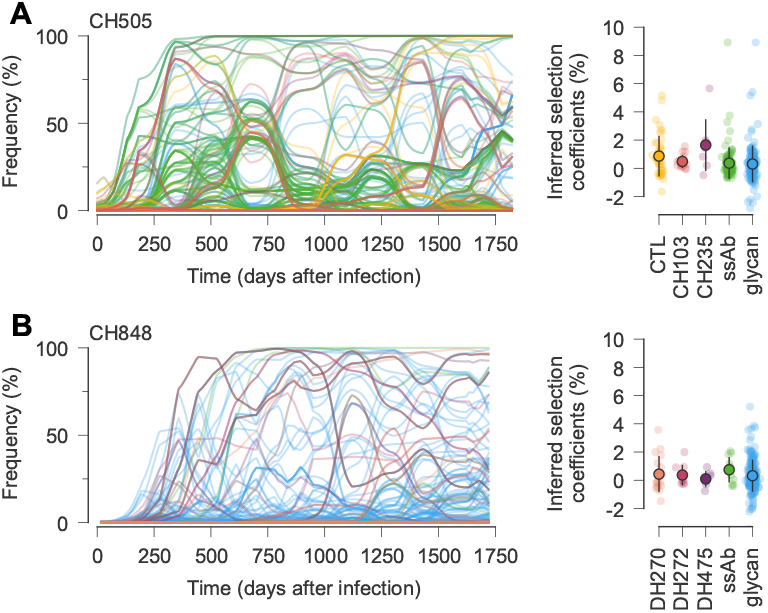
Trajectories and inferred selection coefficients for immune escape mutations and mutations affecting Env glycosylation. **A**, For CH505, early, strongly selected mutations include ones that escape from CD8^+^ T cells and autologous strain-specific antibodies (ssAbs). More moderately selected bnAb resistance mutations tend to arise later. Note that mutations that affect bnAb resistance can appear in the viral population before bnAbs are generated. Open circles and error bars reflect the mean and standard deviation of inferred selection coefficients in each category. **B**, For CH848, mutations affecting glycosylation dominate the early phase of evolution, followed later by mutations affecting bnAb resistance. For easier visualization, frequency trajectories are shown with exponential smoothing with a time scale of *t* = 50 days. See **Supplementary Fig. 3** for a detailed view of mutation frequency trajectories by mutation type without smoothing.

### Consistent patterns of selection in SHIV evolution

Simian-human immunodeficiency viruses (SHIVs) have numerous applications in HIV/AIDS research ^61,62^. Recently, Roark and collaborators studied SHIV-antibody coevolution in rhesus macaques (RMs), which they compared with patterns of HIV-1 evolution ^20^. Two of the SHIV constructs in this study included envelope sequences derived from CH505 and CH848 transmitted/founder (TF) viruses. There, it was found that 2 out of 10 RMs inoculated with SHIV.CH505 and 2 out of 6 RMs inoculated with SHIV.CH848 developed antibodies with substantial breadth.

To understand whether the patterns of HIV-1 selection observed in CH505 and CH848 are repeatable, and to search for viral factors that distinguish between individuals who develop bnAbs and those who do not, we analyzed SHIV.CH505 and SHIV.CH848 evolution in RMs ^20^. To prevent spurious inferences, we first omitted data from RMs with <3 sampling times or <4 sequences in total (Methods). After processing, we examined evolutionary data from 7 RMs inoculated with SHIV.CH505 and 6 RMs inoculated with SHIV.CH848 (**Supplementary Tables 6-7**). We then computed selection coefficients for SHIV mutations within each RM. Reasoning that selective pressures across SHIV.CH505 and SHIV.CH848 viruses are likely to be similar, we also inferred two sets of joint selection coefficients that best describe SHIV evolution in SHIV.CH505- and SHIV.CH848-inoculated RMs, respectively (Methods).

As before, we examined the top 2% of SHIV.CH505 and SHIV.CH848 mutations that we inferred to be the most beneficial for the virus. Overall, we found consistent selection for reversions (17.9- and 14.2-fold enrichment, *p* = 4.3 *×* 10^−11^ and 1.2 *×* 10^−11^ for SHIV.CH505 and SHIV.CH848, respectively), with slightly attenuated enrichment in mutations that affect N-linked glycosylation (2.7- and 4.2-fold enrichment, *p* = 5.4 *×* 10^−2^ and 3.4 *×* 10^−6^). However, there is a small subset of mutations that *shift* glycosylation sites by simultaneously disrupting one N-linked glycosylation motif and completing another, where highly beneficial mutations occur far more often than expected by chance (158.3- and 191.1-fold enrichment, *p* = 1.7 *×* 10^−4^ and 7.9 *×* 10^−11^ for CH505 and SHIV.CH505, respectively; 118.7-fold and 90.4-fold enrichment, *p* = 2.0 *×* 10^−4^ and 2.3 *×* 10^−7^ for CH848 and SHIV.CH848).

Intuitively, one may expect that strongly beneficial SHIV mutations are more likely to be observed in samples from multiple RMs. However, we found that the number of RMs in which a mutation is observed is only weakly associated with the fitness effect of the mutation (**Supplementary Fig. 4**).

While substantially deleterious SHIV mutations are rarely observed across multiple RMs, neutral and nearly neutral mutations are common. Thus, in this data set, it is not generally true that SHIV mutations observed in multiple hosts must significantly increase viral fitness.

We observed some differences between HIV-1 and SHIV in the precise locations of the most beneficial Env mutations. For example, mutations in V4 are highly enriched in CH505 due to a CD8^+^ T cell epitope in this region, but not in SHIV.CH505 (2.7-fold enrichment, *p* = 0.13). For SHIV.CH848, beneficial mutations are modestly enriched in V1 and V5 (4.7- and 4.2-fold, *p* = 4.0 *×* 10^−3^ and 6.7 *×* 10^−2^), as in CH848, but not for V3 (0.9-fold enrichment, *p* = 0.22).

Despite some differences in the top mutations, patterns of selection over time in SHIV were very similar to those found for HIV-1. As before, highly beneficial mutations, including ones affecting glycosylation, tended to appear earlier in infection. This was followed by modestly beneficial mutations at later times, including ones involved in resistance to bnAbs in the RMs who developed antibodies with significant breadth (examples in **Fig. 4**).

**Fig. 4.**
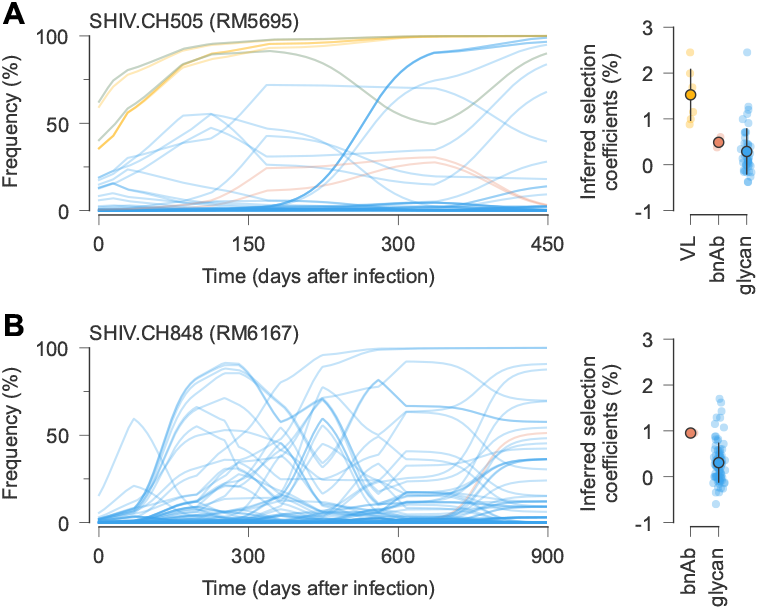
Example trajectories and selection coefficients for SHIV mutations that affect viral load, bnAb recognition, or glycosylation. **A**, In RM5695, infected with SHIV.CH505, mutations known to increase viral load ^60^ and ones affecting glycosylation were rapidly selected. Mutations affecting resistance to broad antibodies ^20^ arose later under moderate selection. Open circles and error bars reflect the mean and standard deviation of inferred selection coefficients in each category. **B**, Slower but qualitatively similar evolutionary patterns were observed in RM6163, infected with SHIV.CH848. For easier visualization, frequency trajectories are shown with exponential smoothing with a time scale of *t* = 50 days.

### Detection of SHIV mutations that increase viral load

A major goal of nonhuman primate studies with SHIV is to faithfully recover important aspects of HIV-1 infection in humans. However, due to the divergence of simian immunodeficiency viruses and HIV-1, SHIVs are not always well-adapted to replication in RMs ^60^. To combat this problem, a recent study identified six SHIV.CH505 mutations that increase viral load (VL) in RMs ^60^. These mutations result in viral kinetics that better mimic HIV-1 infection in humans.

Our analysis readily identifies the SHIV.CH505 mutations shown to increase VL. Five out of the top six SHIV.CH505 mutations with the largest average selection coefficients are associated with increased VL (**Supplementary Table 8**). The final mutation identified by Bauer et al., N130D, is ranked tenth. We also find highly beneficial mutations in SHIV.CH848 that are distinct from those in SHIV.CH505 (**Supplementary Table 9**). Highly-ranked mutations identified here may be good experimental targets for future studies aimed at increasing SHIV.CH848 replication *in vivo*.

### Fitness agreement between HIV-1 and SHIV

Next, we explored the similarity in the overall viral fitness landscapes inferred for HIV-1 and SHIV, beyond just the top mutations. First, we computed the fitness of each SHIV sequence using the joint SHIV.CH505 and SHIV.CH848 selection coefficients inferred from RM data. Then, we computed fitness values for SHIV sequences using selection coefficients inferred from HIV-1 evolution in CH505 and CH848.

We observed a remarkable agreement between SHIV fitness values computed from these two sources (**Fig. 5**). For both SHIV.CH505 and SHIV.CH848, the correlation between viral fitness estimated using data from humans (i.e., CH505 and CH848) and RMs is strongly and linearly correlated (Pearson’s *r* = 0.96 and 0.95, *p <* 10^−20^). This implies that evolutionary pressures on the envelope protein SHIV-infected RMs are highly similar to those on HIV-1 in humans with the same TF Env. In fact, this relationship holds even beyond the SHIV sequences observed during infection. Fitness estimates for sequences with randomly shuffled sets of mutations are also strongly correlated (**Supplementary Fig. 5**). In contrast, the inferred fitness landscapes of CH505 and CH848, which share few mutations in common, are poorly correlated (**Supplementary Fig. 6**). This suggests that the similarities between viral fitness values in humans and RMs are not artifacts of the model, but rather stem from similarities in underlying evolutionary drivers.

**Fig. 5.**
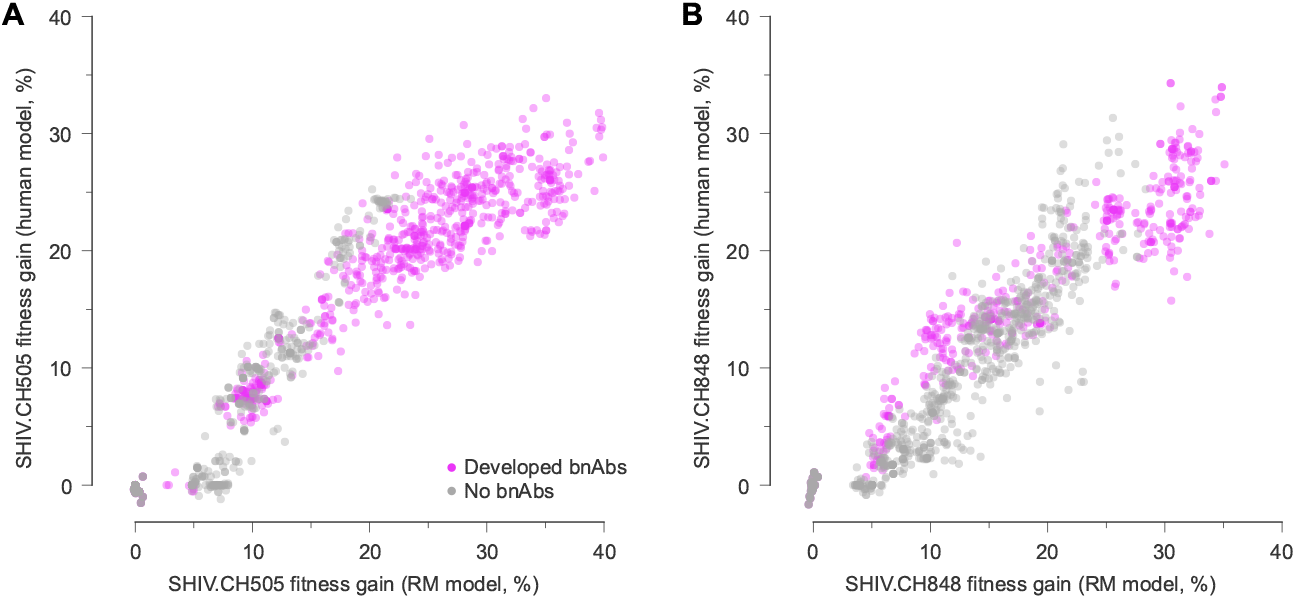
Inferred fitness landscapes in HIV-1 and SHIV are highly similar. **A**, Fitness of SHIV.CH505 sequences relative to the TF sequence across 7 RMs, including 2 that developed bnAbs, 1 that developed tier 2 nAbs that lacked a critical mutation for breadth, and 4 that did not develop broad antibody responses, evaluated using fitness effects of mutations using data from CH505 and using RM data. The fitness values are strongly correlated, indicating the similarity of Env fitness landscapes inferred using HIV-1 or SHIV data. Values are normalized such that the fitness gain of the TF sequence is zero. **B**, Fitness values for SHIV.CH848 sequences also show strong agreement between CH848 and SHIV.CH848 landscapes.

### Evolutionary dynamics forecast antibody breadth

Given the similarity of HIV-1 and SHIV evolution, we sought to identify evolutionary features that distinguish between hosts who develop broad antibody responses and those who do not. **Figure 5** shows that SHIV sequences from hosts with bnAbs often reach higher fitness values than those in hosts with only narrow-spectrum antibodies. We hypothesized that stronger selective pressures on the virus might drive viral diversification, stimulating the development of antibody breadth. Past studies have associated higher viral loads with bnAb development and observed viral diversification around the time of bnAb emergence ^13,16,17^. Computational studies and experiments have also shown that sequential exposure to diverse antigens can induce cross-reactive antibodies ^63–65^.

To further quantify SHIV evolutionary dynamics, we computed the average fitness gain of viral populations in each RM over time. We observed a striking difference in SHIV fitness gains between RMs that developed broad antibody responses and those that did not (**Fig. 6**). In particular, SHIV fitness increased rapidly before the development of antibody breadth. SHIV fitness gains in RM5695, which developed exceptionally broad and potent antibodies ^20^, were especially rapid and dramatic. These fitness differences were not attributable to bnAb resistance mutations, which were only moderately selected and generally appeared after bnAbs developed.

**Fig. 6.**
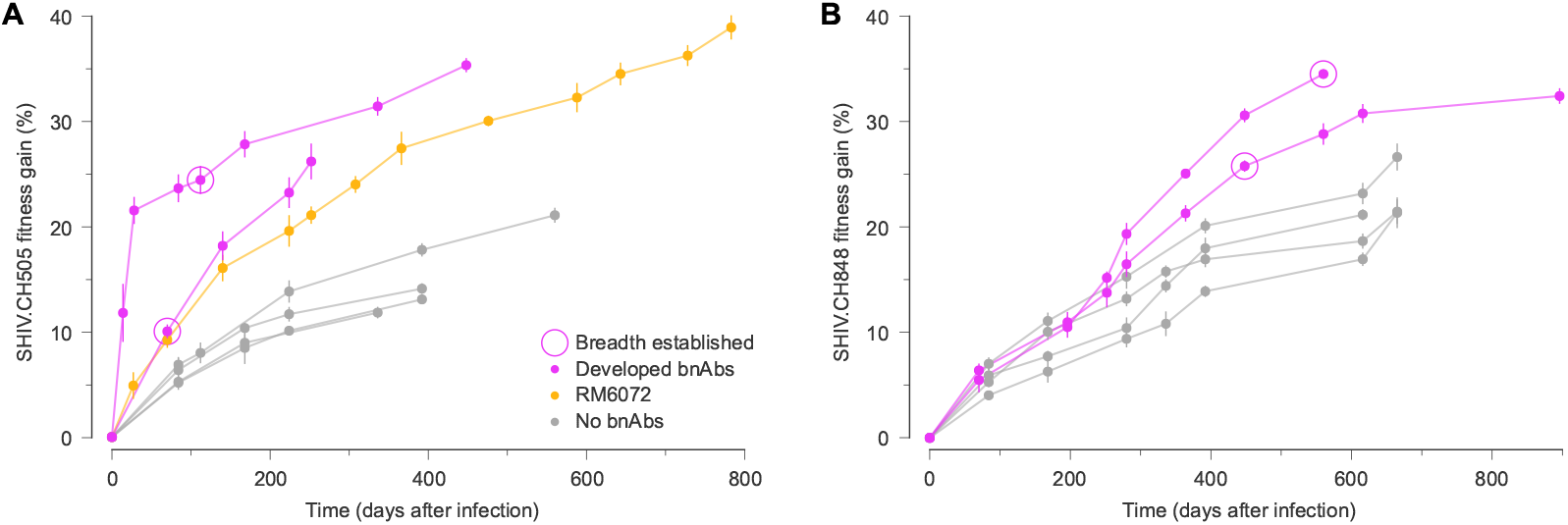
Rapid SHIV fitness gains precede the development of broadly neutralizing antibodies. For both SHIV.CH505 (**A**) and SHIV.CH848 (**B**), viral fitness gains over time display distinct patterns in RM hosts that developed bnAbs versus those that did not. Notably, the differences in SHIV fitness gains between hosts with and without broad antibody responses appear *before* the development of antibody breadth, and cannot be attributed to selection for bnAb resistance mutations. RM6072 is an unusual case, exhibiting antibody development that was highly similar to CH505. Although RM6072 developed tier 2 nAbs, they lacked key mutations critical for breadth ^20^. Points and error bars show the mean and standard deviation of fitness gains across SHIV samples in each RM at each time.

One outlier in this pattern is RM6072, infected with SHIV.CH505. Antibody development in RM6072 followed a path that was remarkably similar to CH505, including a lineage of antibodies, DH650, directed toward the CD4 binding site of Env ^20^. However, resistance to the DH650 lineage is conferred by a strongly selected mutation that adds a glycan at site 234 (T234N, with an inferred selection coefficient of 4.5%). Broadly neutralizing antibodies similar to DH650 are able to accommodate this glycan due to shorter and/or more flexible light chains ^20,66^, but DH650 cannot. Antibody evolution in RM6072 thus proceeded along a clear pathway toward bnAb development, but lacked critical mutations to achieve breadth.

Next, we quantified how different types of SHIV mutations contributed to viral fitness gains over time. We examined contributions from VL-enhancing mutations ^60^, antibody escape mutations ^20,60^, other mutations affecting Env glycosylation, and reversions to subtype consensus. We found increased fitness gains across all types of mutations in RMs that developed broad antibody responses, compared to those that did not (**Supplementary Fig. 7**). VL-enhancing mutations, known antibody resistance mutations, and reversions typically made the largest contributions to viral fitness.

### Robustness of inferred selection to changes in the fitness model and finite sampling

In the analysis above, we used a simple model where the net fitness effect of multiple mutations is simply equal to the sum of their individual effects. Recently, methods have also been developed that can infer epistatic fitness effects from data, which include pairwise interactions between mutations ^40,41^. We reanalyzed these data to examine how inferred fitness changes when epistasis is included in the model, using the approach of Shimagaki and Barton ^41^ (Methods). Overall, the inferred epistatic interactions were modest (**Supplementary Fig. 8**). In CH505, we found that the CD4 binding site, V1 (especially sites 136–146 in HXB2 numbering) and V5 regions were modestly but significantly enriched in the most beneficial (top 1%) of epistatic interactions (2.5-, 1.2- and 1.8-fold enrichment with *p* = 1.0 *×* 10^−21^, 6.3 *×* 10^−6^ and 6.3 *×* 10^−5^, respectively). Epistatic interactions between N280S/V281A and E275K/V281G, which confer resistance to CH235 (ref. ^22^), ranked in the top 6.5% and 13.0% of interactions. In CH848, we found 1.3-, 1.5-, and 2.3-fold enrichment in strong beneficial epistatic interactions in the CD4 binding site, V4, and V5 regions, respectively (*p* = 4.0 *×* 10^−6^, *p* = 2.5 *×* 10^−14^ and 3.2 *×* 10^−19^).

To compare the typical fitness effects of individual mutations in the model with epistasis to those in the additive model, we computed effective selection coefficients for the epistatic model. For each mutant allele *a* at each site *i*, we computed the average difference in fitness between sequences in the data set with the mutation and hypothetical sequences that are the same as those in the data, except with the mutant allele *a* reverted to the TF one. In this way, the effective selection coefficient measures the typical effect of each mutation in the data set, while also accounting for epistatic interactions with the sequence background. We found that the effective selection coefficients were highly correlated with the selection coefficients from the additive model (**Supplementary Fig. 9**). We also found strong agreement between the additive and epistatic model fitness values for each sequence in both HIV-1 and SHIV data (**Supplementary Fig. 10**).

Finite sampling of sequence data could also affect our analyses. To further test the robustness of our results, we inferred selection coefficients using bootstrap resampling, where we resample sequences from the original ensemble, maintaining the same number of sequences for each time point and subject. The selection coefficients from the bootstrap samples are consistent with the original data (see **Supplementary Fig. 11** for a typical example), with Pearson’s *r* values of around 0.85 for HIV-1 data sets and 0.95 for SHIV data sets, respectively.

## Discussion

HIV-1 evolves under complex selective pressures within individual hosts, balancing replicative efficiency with immune evasion. Here, we quantitatively studied the evolution of HIV-1 and SHIV (featuring HIV-1-derived Env sequences) across multiple hosts, including some who developed broad antibody responses against the virus. Our study highlighted how different classes of mutations (e.g., mutations affecting T cell escape or Env glycosylation) affect fitness *in vivo*. In both HIV-1 and SHIV, we found strong selection for reversions to subtype consensus and some mutations that affected N-linked glycosylation motifs or resistance to autologous strain-specific antibodies. Few CD8^+^ T cell epitopes were identified in this data set, but the T cell escape mutations that we did observe were highly beneficial for the virus. Consistent with past work studying VRC26 escape in CAP256, we observed more modest selection for bnAb resistance mutations ^39^.

Overall, we found striking similarities between Env evolution in humans and RMs. Importantly, these parallels extend beyond the observation of repetitive mutations: the number of hosts in which a mutation was observed was only weakly associated with the mutation’s fitness effect (**Supplementary Fig. 4**). Our inferred Env fitness values in humans and RMs were highly correlated, indicating that the functional and immune constraints shaping Env evolution in HIV-1 and SHIV infection are very similar. Our findings therefore reinforce SHIV as a model system that closely mirrors HIV-1 infection.

We discovered that the speed of SHIV fitness gains was clearly higher in RMs that developed broad antibody responses than in those with narrow-spectrum antibodies. Fitness gains in the viral population preceded the development of bnAbs, and they were not driven by bnAb resistance mutations. This suggests that rapid changes in the viral population are a cause rather than a consequence of antibody breadth. While our sample is limited to 13 RMs and two founder Env sequences, we find a clear separation between RMs that did or did not develop antibody breadth. Thus, the dynamics of viral fitness may serve as a quantitative signal associated with bnAb development.

The induction of bnAbs is a major goal of HIV-1 vaccine design ^8^. Both computational ^63,65,67,68^ and experimental ^64,69,70^ studies, as well as observations from individuals who developed bnAbs ^13,14,22^, suggest that the co-evolution of antibodies and HIV-1 is important to stimulate broad antibody responses. Our results could thus inform HIV-1 vaccine research. While precise immune responses and viral escape pathways can differ across individuals, the quantitative similarity in viral evolutionary constraints across humans and RMs suggests that SHIV data can provide a valuable source of information about Env variants that contribute to bnAb development, especially when detailed longitudinal data from humans does not exist. While the concept of sequential immunization is well-established ^10,63,64,71,72^, our findings also suggest a possible new design principle. Immunogens could be engineered to reproduce the dynamics of viral population change that are associated with rapid fitness gains, which we found to precede the emergence of bnAbs. This emphasis on broader, population-level dynamics could complement investigations of the molecular details of virus and antibody coevolution.

As noted above, Roark and collaborators also performed a detailed comparison of HIV-1 and SHIV evolution with the same TF Env sequences ^20^. One of their main conclusions was that most Env mutations were selected for escape from CD8^+^ T cells or antibodies. We found that many antibody resistance mutations identified by Roark et al. are also positively selected in our analysis. Mutations at sites 166 and 169 were shown to confer resistance to a V2 apex bnAb, RHA1, isolated in RM5695 (ref. ^20^). We inferred moderately positive selection coefficients of 0.49% and 0.43% for R166K and R169K, respectively. The same mutations were found in RM6070, which also developed V2 apex bnAbs, with a selective advantage of 1.7% (**Supplementary Table 10**). Mutations conferring resistance to autologous strain-specific nAbs were identified at multiple sites by Roark and colleagues: 130, 234, 279, 281, 302, 330, and 334 in RM6072, which developed antibody responses targeting the CD4 binding site (DH650) and V3 (DH647 and DH648) regions. Mutations Y330H and N334S, which confer resistance to V3 autologous nAbs, were detected in all RMs infected with SHIV.CH505, with selective advantages of 3.0% and 4.6% in RM6072, and 1.7% and 3.2% on average across RMs, respectively. Overall, we found that mutations conferring resistance to autologous strain-specific antibodies were common and more strongly selected than bnAb resistance mutations (**Supplementary Tables 10-11**).

We note that our conclusions about the phenotypic effects of HIV-1 mutations under selection are constrained by the available data. While we observed strong selection for strain-specific antibody resistance mutations, these results could also be affected by the effects of these mutations on viral replication independent of immune escape. In particular, many ssAb resistance mutations are also reversions to the subtype consensus sequence, which have often been observed to improve viral fitness ^39,54^. For example, N334S, K302N, and T234N are all reversions. These are among the most beneficial mutations inferred for SHIV.CH505 (**Supplementary Table 8**). In future work, it would be interesting to attempt to fully separate the fitness effects of mutations due to antibody escape and intrinsic replication (cf. ^42^). Although we have systematically compiled information about mutations known to affect antibody resistance and glycosylation, this data is necessarily incomplete. Some of the strongly beneficial mutations with unknown functional effects that we observe could therefore reflect escape from unmapped immune responses.

There are additional methodological and technical limitations that should be considered in the interpretation of our results. Most notably, we assume that the viral fitness landscape is static in time. While we do not expect selection for effective replication (“intrinsic” fitness) to change substantially over time, pressure for immune escape could vary along with the immune responses that drive them. In prior work, we have found that constant selection coefficients typically reflect the average fitness effect of a mutation when its true contribution to fitness is time-varying ^42,43^. This may not adequately description mutational effects that undergo large or rapid shifts in time. Future work should also examine temporal patterns in selection for individual mutations.

While we found a strong relationship between viral fitness dynamics and the emergence of bnAbs, it may not be true that the former stimulates the latter. For example, bnAbs may have been present within each host before they were experimentally detected. Rapid viral fitness gains within hosts that developed broad antibody responses could then have been driven by undetected bnAb lineages. However, we did not find strong selection for known bnAb resistance mutations, and in at least one case (RM5695), rapid fitness gains (roughly 2 weeks after infection) substantially preceded bnAb detection (16 weeks). Still, given the limited size of the data set that we studied, it is unclear the extent to which our results will transfer to larger and broader data sets.

Among other analyses, Roark et al. used LASSIE ^73^ to identify putative sites under selection (**Supplementary Tables 12-13**). This method works by identifying sites where non-TF alleles reach high frequencies. We found modest overlap between the sites under selection as identified by LASSIE and the mutations that we inferred to be the most strongly selected. For SHIV.CH505, the E640D mutation at site 640 identified by LASSIE is ranked second among 664 mutations in our analysis, and mutations at the remaining 5 sites identified by LASSIE are all within the top 20% of mutations that we infer to be the most beneficial. For SHIV.CH848, the R363Q mutation that is ranked first in our analysis appears at one of the 17 sites identified by LASSIE. Some mutations at the majority of these 17 sites fall within the top 20% most beneficial mutations in our analysis, but some are outliers. In particular, we infer both S291A/P to be somewhat deleterious, with S291P ranked 810th out of 863 mutations.

Beyond the specific context of HIV-1 and bnAb development, our study also provides insight into viral evolution across hosts and related host species. Parallels between the HIV-1 and SHIV fitness landscapes that we infer suggest that there are strong constraints on viral protein function, with few paths to significantly higher fitness. This is consistent with the ideas of methods that use sequence statistics across multiple individuals and hosts to predict the fitness effects of mutations ^74–80^. However, the relationship between the number of individuals in which a mutation was observed and its inferred fitness effect was fairly weak. This suggests that mutational biases and/or sequence space accessibility may play significant roles in short-term viral evolution, even for highly mutable viruses such as HIV-1 and SHIV. As described above, high frequency mutations were also not necessarily highly beneficial. While the recombination rate of HIV-1 is high, correlations between mutations persist, making it difficult to unambiguously interpret frequency changes as signs of selection ^39^.

Our results also point to strong similarities in the immune environment across closely related host species, including preferential targeting of specific parts of viral surface proteins by antibodies. This is supported by the enrichment of beneficial mutations within variable loop regions and at sites that affect the glycosylation of Env. However, despite these constraints, there may still exist a large number of neutral or nearly-neutral mutational paths that remain unexplored.

Overall, our findings support the potential predictability of viral evolution, at least over short time scales. While there are contingencies in evolution – for example, disparate host immune responses or strong epistatic constraints between mutations – these are not so pervasive that they completely change the effective viral fitness landscape or paths of evolution across hosts, given the same founder virus sequence. Similar observations of parallel evolution in HIV-1 have been reported in monozygotic twins infected by the same founder virus ^81^, common patterns of immune escape across hosts ^78,82^ and drug resistance ^83–85^, and long-term experimental evolution ^86^. Our results thus contribute to a growing body of research identifying predictable features in viral evolution. Understanding such features could ultimately inform practical applications such as anticipating the emergence of drug resistance or designing vaccines to limit likely pathways of escape.

## ACKNOWLEDGEMENTS

The work of K.S.S. and J.P.B. reported in this publication was supported by the National Institute of General Medical Sciences of the National Institutes of Health under Award Number R35GM138233.

## AUTHOR CONTRIBUTIONS

All authors contributed to the research design, methods development, interpretation of results, and writing the paper. K.S.S. performed simulations and computational analyses. J.P.B. supervised the project.

## Methods

### Data

We retrieved HIV-1 sequences from CH505 (703010505) and CH848 (703010848) from the HIV sequence database at Los Alamos National Laboratory (LANL) ^87^. The rhesus macaque (RM) SHIV sequences ^88^ were obtained from GenBank ^89^. We then co-aligned SHIV.CH505 and SHIV.CH848 sequences with CH505 and CH848 HIV-1 sequences, respectively, using HIValign ^90^.

#### CH505

CH505 developed two distinct lineages of CD4 binding site (CD4bs) bnAbs, CH103 and CH235 (refs. ^91,92^). CH103 antibodies were detectable by 14 weeks after infection and further developed neutralization breadth between 41-92 weeks ^93^. IC50 values of CH235 against the TF virus were 6.5-fold lower than those of CH103 (ref. ^91^). CH235 lineages could neutralize autologous viruses at week 30. However, viruses that acquired mutations at loop D from 53-100 weeks escaped CH235 neutralization ^91^. Although the neutralization breadth of CH235 was not as broad as that of CH103, this lineage played a critical role; escaping mutations from the CH235 lineage stimulated the development of another lineage with broader neutralization depth ^91^. Mutations in loop D enabled the virus to escape from CH235, but these sequential mutations in loop D, such as E275K, N279D, and V281S, favorably bound to the mature CH103 and continuously increased the binding affinity between mature CH103 and loop D ^91^. Gradually, CH103 matured, developing a broader neutralization breadth.

#### CH848

CH848 developed DH270, a bnAb that targets the glycosylated site adjacent to the third variable loop (V3). DH270 was detectable three and a half years after infection ^94^. Similar to the CH505 case, the CH848 case exhibited cooperative virus and antibody coevolution. The earlier antibody lineages, DH272 and DH475, could neutralize autologous viruses until week 51 and weeks 15-39, respectively. The virus escaped from DH272 and DH475 afterward, with escape mutations including a longer V1V2 loop. DH270 then developed, with potent and broad neutralization breadth ^94^.

#### Rhesus macaques

Chimeric viruses, SHIVs, were constructed by bearing the transmitted/founder (TF) Env from three HIV-1 patients, including CH505 and CH848 (ref. ^88^). In some RMs, SHIV developed similar patterns of mutations to those observed in human donors. In our analysis, we considered RMs with SHIV sequences sampled at at least three points in time. This yielded a set of 7 RMs and 6 RMs for SHIV.CH505 and SHIV.CH848, respectively. The **Supplementary Tables 6-7** summarize the number of sequences, time points, and the development of bnAbs for each individual in the SHIV cases as well as HIV-1 cases.

### Sequence data processing

#### Data quality control

To focus our analysis on functional sequences, we removed sequences with more than 200 gaps. To eliminate rare insertions or possible alignment errors, we also masked sites where gaps occurred in more than 95% of sequences within each individual host. To limit errors in virus frequencies, we only considered data from time points with four or more sequences.

#### Identifying reversions

A mutation is classified as a reversion if the new (mutant) nucleotide matches with the nucleotide at the same site in the HIV-1 consensus sequence from the same subtype. Here, all viruses were subtype C, so we compared with the subtype C consensus sequence as defined by LANL.

#### Identifying mutations that affect N-linked glycosylation

To identify mutations that affect glycosylation, we search for Env mutations that modify the N-linked glycosylation motif Asn-X-Ser/Thr, where X can be any amino acid except proline. We identified three types of mutations affecting glycosylation: “shields,” which complete a previously incomplete glycosylation motif, “holes,” which disrupt an existing glycosylation motif, and “shifts,” which simultaneously complete one N-linked glycosylation motif and disrupt another.

#### Enrichment analysis

We used fold enrichment values and Fisher’s exact test to quantify the excess or lack of mutations. For a particular subset of mutations (for example, the top *x*% beneficial mutations), we first computed the number of mutations in that subset that do (*n*_sel_) and do not (*N*_sel_) have a particular property (for example, nonsynonymous mutations in the CD4 binding site). We then computed the total number of mutations that do and do not have the property (*n*_null_ and *N*_null_, respectively) across the entire data set. The fold enrichment value is then 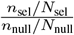. The term 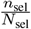 quantifies the fraction of mutations having specific properties across the selected mutations, while the denominator 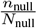 is the fraction of all mutations that have the property. Fisher’s exact *p* values are computed from the 2 *×* 2 table with *n*_sel_, *n*_null_ *n*_sel_ in the first row, and *N*_sel_ − *n*_sel_, *N*_null_ − *N*_sel_ − (*n*_null_ − *n*_sel_) in the second row ^95^.

### Inferring fitness effects of mutations

In this section, we describe the inference framework used to infer the fitness effects of mutations (selection coefficients) from temporal genetic data.

#### Evolutionary model

We model viral evolution with the Wright-Fisher (WF) model, a fundamental model in population genetics ^96^. In this model, a population of *N* individuals (viruses or infected cells, in our case) undergo discrete rounds of selection, mutation, and replication. Each genotype *α* is represented by a sequence 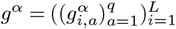, where 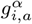 is equal to one if genotype *α* has allele *a* at locus *i* and zero otherwise. Here *L* and *q* represent the length of the genetic sequence (number of loci) and the number of statues at each locus (i.e., number of nucleotides or amino acids), respectively. We use *q* = 5 and *q* = 21 for DNA and amino acid sequences, respectively, in real HIV-1 and SHIV data.

We define the fitness of an individual with genetic sequence *g* by

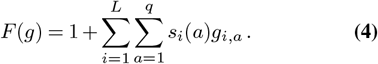

Here *s*_*i*_(*a*) is a selection coefficient, quantifying the fitness effect of allele *a* at locus *i*. If *s*_*i*_(*a*) *>* 0, the allele *a* is beneficial (enhancing replication), and if *s*_*i*_(*a*) *<* 0 it is deleterious (impairing replication). By convention, we set the selection coefficient for TF alleles to zero. Individuals with higher fitness values are more likely to replicate than those with lower fitness.

Mutations introduce new genotypes and drive the evolution of the population. Let us define *µ*^*αβ*^ as the probability of mutation from genotype *α* to genotype *β* per replication cycle. Below, we will express this probability in terms of a mutation rate per site per round of replication. In the analysis of real data, we use asymmetric mutation rates estimated from intra-host HIV-1 data ^97^.

Given these parameters, the WF model describes the dynamics of the frequencies of different genotypes in the population over time. We write the frequency of genotype *α* at time *t* as *z*_*α*_(*t*). Given that the frequency of genotypes in the population at time *t* is *z*(*t*) = (*z*_1_(*t*), *z*_2_(*t*), …, *z*_*M*_ (*t*)), where *M* is the total number of genotypes, the probability distribution of the frequency of genotypes in the next generation *z*(*t* + 1) is

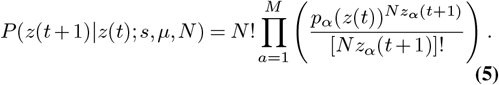

Here *p*_*α*_ is

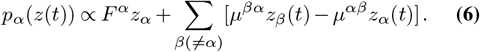

where *F*^*α*^ is the fitness value of genotype *α*, based on Eq. (4). Across *K* generations, the probability of an entire evolutionary trajectory, defined by the vector of genotype frequencies at each time, is then

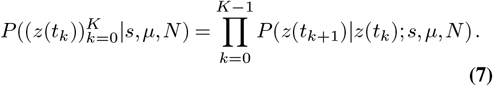

#### Diffusion limit

When the population size is sufficiently large, the evolution of the population defined in Eq. (5) can be reasonably well approximated by a Gaussian process, which is a solution to the Fokker-Planck (forward Kolmogorov) equation ^98,99^.

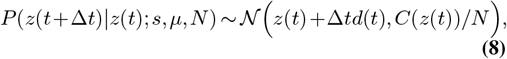

with the drift vector *d*(*t*) and the diffusion matrix *C*(*z*(*t*))*/N* such that

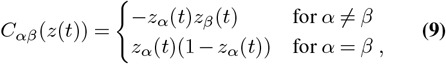

and

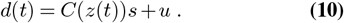

#### Dimensional reduction

While the WF process in genotype space provides valuable insights into genotype dynamics, the mathematical expressions are sometimes challenging to interpret. To obtain more intuitive expressions, we can project the dynamics onto the space of allele frequencies,

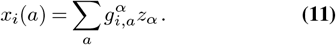

One can then find the drift vector *d* and diffusion matrix *C/N* in allele frequency space,

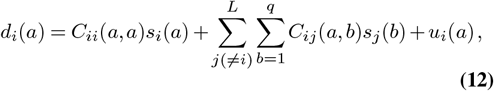

and

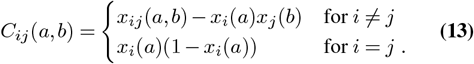

Here, *x*_*ij*_(*a, b*) is the frequency of individuals with alleles *a* and *b* at loci *i* and *j*, and *u*_*i*_(*a*) is net expected change in frequency of allele *a* at *i* due to mutations, which is given explicitly in Eq. (17) below. The first term in Eq. (**??**) gives the expected change in the frequency *x*_*i*_(*a*) due to the direct fitness effect *s*_*i*_(*a*), while the second term represents the contributions due to indirect or genetic linkage effects with other alleles *j*.

#### Maximum path likelihood

Following recent work ^100^, we employed Bayes’ rule to find the selection coefficients that best explain the data. These are the coefficients that maximize the posterior distribution

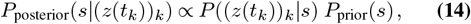

which is a product of the likelihood of the evolutionary trajectory observed in the data Eq. (7) (under the diffusion limit Eq. (8)) and a prior distribution for the selection coefficients. We chose a Gaussian prior distribution with zero mean and a covariance of *I/*(*Nγ*), where *I* is the identity matrix. This prior distribution penalizes the inferences of large selection coefficients when they are not well-supported by the data. The maximum *a posteriori* selection coefficients are then given by ^100^

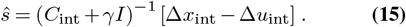

Here *C*_int_, Δ*x*_int_, and Δ*u*_int_ represent the covariance matrix, vector of frequency changes, and mutational flux integrated over the evolution

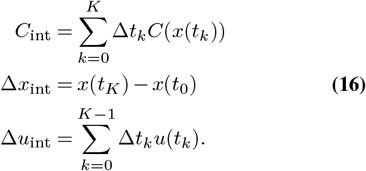

The mutational flux *u* is characterized by the rates of mutations from nucleotides *b* to *a*, denoted by *µ*_*ab*_, which are determined from longitudinal HIV-1 populations in untreated patients ^97^. The change of the *a* nucleotide frequency at locus *i* due to mutation is given by:

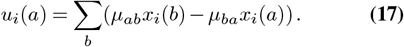

Inverting the integrated covariance matrix effectively reveals the underlying direct allele interactions and resolves the genetic linkage effects.

The shift in the covariance diagonal in Eq. (15), arising from the selection coefficients’ posterior distribution, reflects the uncertainty in the selection distribution. We used *γ* = 10 for all data sets, but the model is robust to variation in the strength of regularization ^100^. For the mutation rates, we incorporated the transition probabilities among arbitrary DNA nucleotides, estimated from whole-genome deep sequencing of multiple untreated HIV-1 patients followed for 5 to 8 years post-infection ^97^.

#### Integration of covariance

When the time interval of the observation Δ*t* is sufficiently short, the trajectory of the allele frequency would be continuous and ideally it would be a smooth curve ^101^. To accurately estimate the covariance matrix, we employ piecewise linear interpolation for frequencies. Let *τ* ∈ [0, 1], then the linear interpolation for a frequency vector can be expressed as:

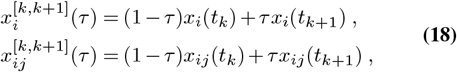

which yields,

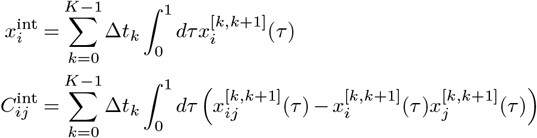

For simplicity in notation, we omitted nucleotide indices. The explicit expression of the integrated covariance is given in refs. ^100,101^.

### Joint RM model

In addition to fitness models derived from SHIV data for individual RMs, we inferred a joint model under the assumption that virus evolution within each individual RM with the same TF virus is governed by a similar fitness landscape. This method improves inference accuracy from the WF process and deep mutational scanning data ^102–104^. The joint path likelihood for allele frequency trajectories across RMs with the same TF virus is then

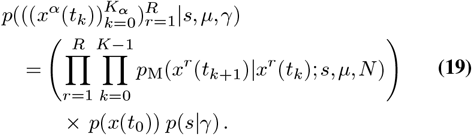

Here, *x*^*r*^ is the allele frequency of the *r*-th individual and *R* is the number of replicate individuals (i.e., the number of RMs sharing the same TF virus). The initial state is *p*(*x*^*r*^(*t*_0_)) = *p*(*x*(*t*_0_)) for all *r* for individuals with the same TF virus. The solution of the joint path likelihood is given by

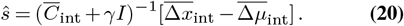

Here, the overbar denotes the sum over the replicate RMs. We emphasize that the joint selection coefficients in Eq. (20) are not the same as selection coefficients that are simply averaged across RMs with the same TF virus. The joint selection coefficients are more robust, as they are guided by the level of evidence within each individual rather than naive averaging.

### Geometrical interpretation of the fitness comparison

The Pearson values we utilized to compare the fitness landscapes denoted as *F*_*s*_(*g*) := *F* (*g* | *s*) and *F*_*h*_(*g*) := *F* (*g* | *h*) can be expressed by the following simple relation:

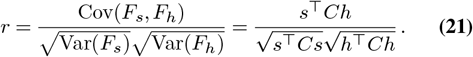

Here, Cov(*F*_*s*_, *F*_*h*_) and Var(*F*_*s*_) represent the covariance and variance values estimated from the samples being compared, (*F*_*s*_(*g*^*n*^), *F*_*h*_(*g*^*n*^))_*n*_. *C* is the covariance matrix defined between arbitrary loci. The last equation can be interpreted as an angle between two vectors, *s* and *h*, with a metric matrix *C*; if *s* = *h*, the Pearson value clearly becomes 1. However, the “similarity” also depends on how these vectors are projected by *C*; eigenmodes associated with larger variance of statistics will be more emphasized. The last expression readily implies an interpretation for the case of shuffled sequences; shuffling the sequences equates to diluting the covariance between loci, resulting in the metric matrix becoming a diagonal matrix. Removing the off-diagonal elements corresponds to lifting the constraints on the fitness landscape.

### Epistatic fitness model

We consider the following pairwise epistatic fitness function, which depends on epistatic interactions *s*_*ij*_ across all possible pairs of loci:

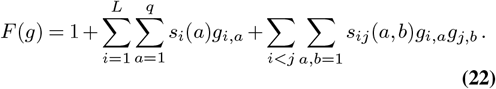

Our goal is to obtain the epistatic interactions *s*_*ij*_(*a, b*) as well as the selection coefficients *s*_*i*_(*a*) from temporal genetic sequences.

The basic logic for inferring these fitness parameters parallels the additive case. The only practical difference is that epistatic interactions can influence the dynamics of additive and pairwise mutation frequencies. Under the diffusion limit, we obtain the following drift terms, which align with Eq. (12) in the additive model,

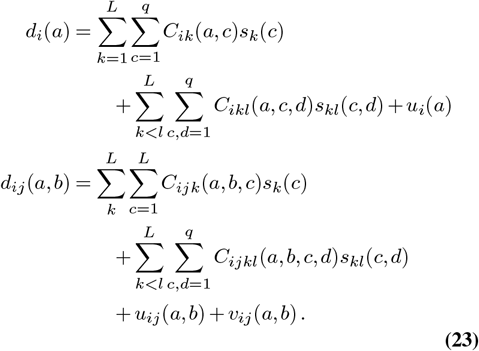

and diffusion matrices,

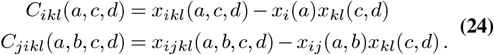

Here, *u*_*i*_ and *u*_*ij*_ represent the expected frequency changes due to mutations for additive and pairwise terms, while *v*_*i*_ represents the changes in pairwise frequencies due to recombination. These explicit expressions indicate that the *u*_*i*_ remains the same as in the additive fitness case; therefore, Eq. (17) holds. The pairwise term is given as

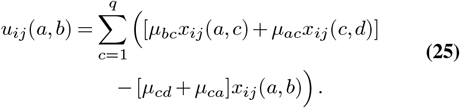

and the *v*_*ij*_ is expressed as

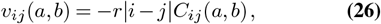

where *r* denotes the recombination rate. In this study, we set *r* = 10^−5^. More detailed derivations are provided in the previous studies ^102,103^.

The technical challenge of the epistasis inference is that the diffusion matrix *C* involves third- and fourth-order interactions, and the number of matrix elements scales as 𝒪 ((*qL*)^4^), while the computational cost to invert it scales as 𝒪 ((*qL*)^6^). Recently, a more efficient computational method was proposed, reducing both the necessary memory usage and computational times by 𝒪 ((*qL*)^2^) (ref. ^103^). The essential idea involves factorizing the higher-order covariance matrix using the rectangular matrix Ξ∈ ℝ^*D×d*^ such that *C* = ΞΞ^*⊤*^, where *D* scales as 𝒪 ((*qL*)^2^) while *d ≪ D*. This method resolves the linear equation without obtaining an explicit expression of *C* and avoids any computations involving more than *O*((*qL*)^2^) operations ^103^.

### Gauge transformation

Since constant shifts in fitness parameters do not affect relative fitness, it is always possible to transform fitness values without changing the resulting genotype distribution. For example, in the additive model, individual selection coefficients can be shifted as

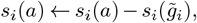

where 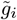 is the allele of a chosen reference genotype 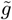 (e.g., the TF sequence) at site *i*. This transformation preserves relative fitness but it ensures that 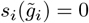, making inferred selection coefficients more interpretable.

In statistical physics, such transformations, where model parameters are changed without altering the underlying probability distribution—are referred to as gauge transformations ^105,106^. Similar transformations have been employed in recent studies to improve interpretability and sparsity in epistatic models ^107^. We can apply an analogous transformation to epistatic interactions:

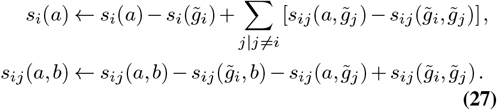

Under this transformation, any selection coefficients or epistatic terms involving TF alleles are zero by definition, while relative fitness remains unchanged.

### Regularization

Regularization is used to reduce the effective number of parameters in the fitness model. In our analysis, we applied strong regularization (*γ* = 10^10^) to any selection or epistatic coefficients involving TF alleles, ensuring they are effectively zero under the gauge transformation. Following prior work ^108^, we penalized epistatic interactions between loci more than 50 nucleotides apart on the reference sequence with the same strong regularization. We also used the same moderate regularization value of *γ* = 50 for all other epistatic terms, and used *γ* = 10 for selection coefficients, consistent with the additive model.

## Code availability

Sets of processed data and computer code used in this study are available in the GitHub repository of https://github.com/bartonlab/paper-HIV-coevolution. This repository contains source files that process HIV-1 and SHIV sequences, infer selection coefficients, and identify and characterize mutations. The included Jupyter notebooks can be run to reproduce the figures presented here. The original HIV-1 sequences can be retrieved from the LANL database (https://www.hiv.lanl.gov/content/index), and SHIV sequences can be found at GenBank (https://www.ncbi.nlm.nih.gov/).

---

**Supplementary Fig. 1.**
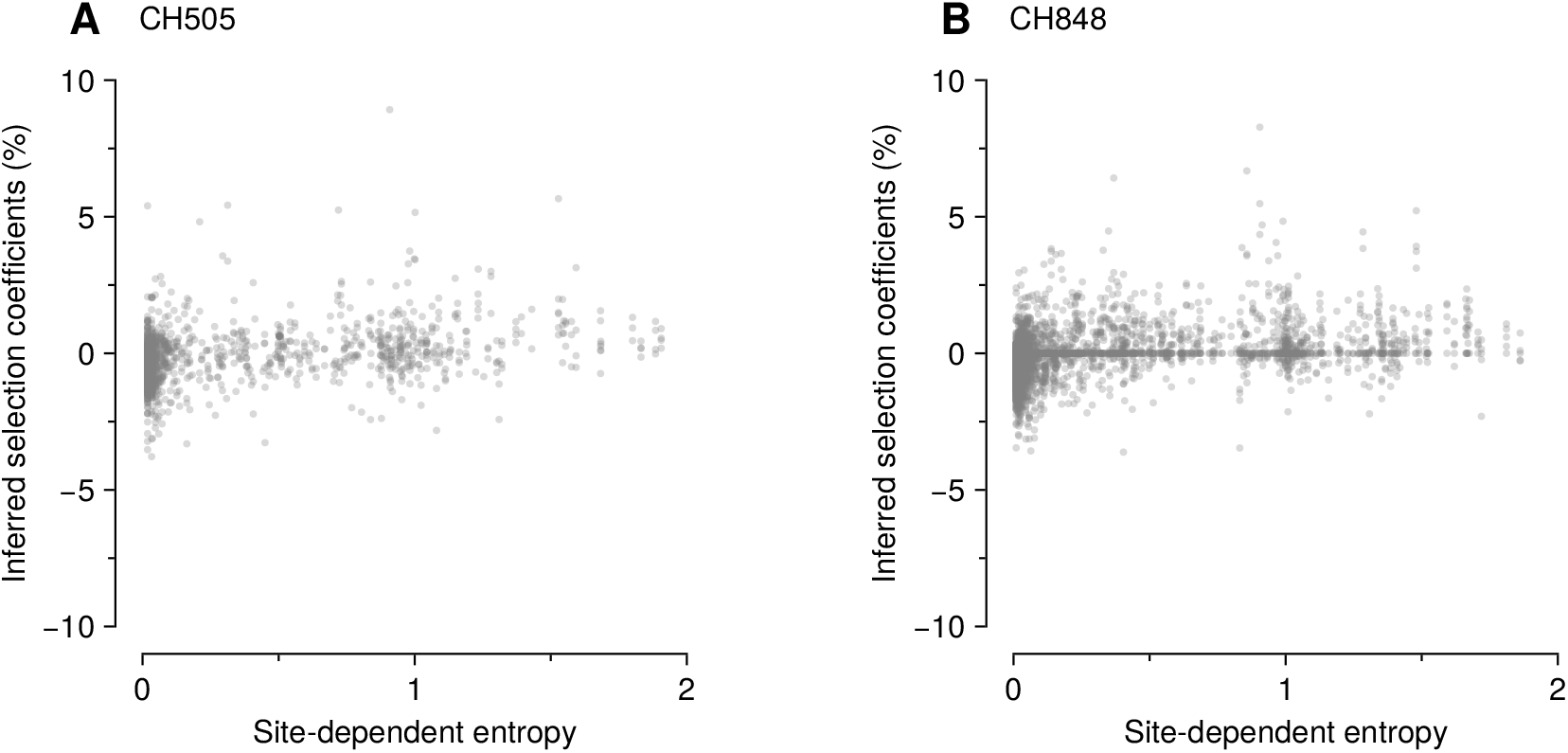
Weak correlation between sequence variability, as measured by entropy, and inferred selection. To quantify sequence variability, we computed “site-dependent entropy” scores 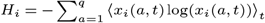, for each site *i* in HIV-1 sequences from CH505 and CH848. In the expression for *H, x* (*a, t*) represents the frequency of allele *a* at site *i* at time *t*, and ⟨*·*⟩_*t*_ denotes the average over time points. For both CH505 (**A**) and CH848 (**B**), we found no clear systematic association between the entropy values and beneficial inferred selection coefficients. Pearson’s *r* values between entropy and inferred selection coefficients are 0.32 and 0.30 for CH505 and CH848, respectively. In particular, large selection coefficients are not concentrated among sites with the highest entropy values. Among the mutations in the top 5% inferred selection coefficients, the *r* values are −0.01 and 0.11 for CH505 and CH848, respectively.

**Supplementary Fig. 2.**
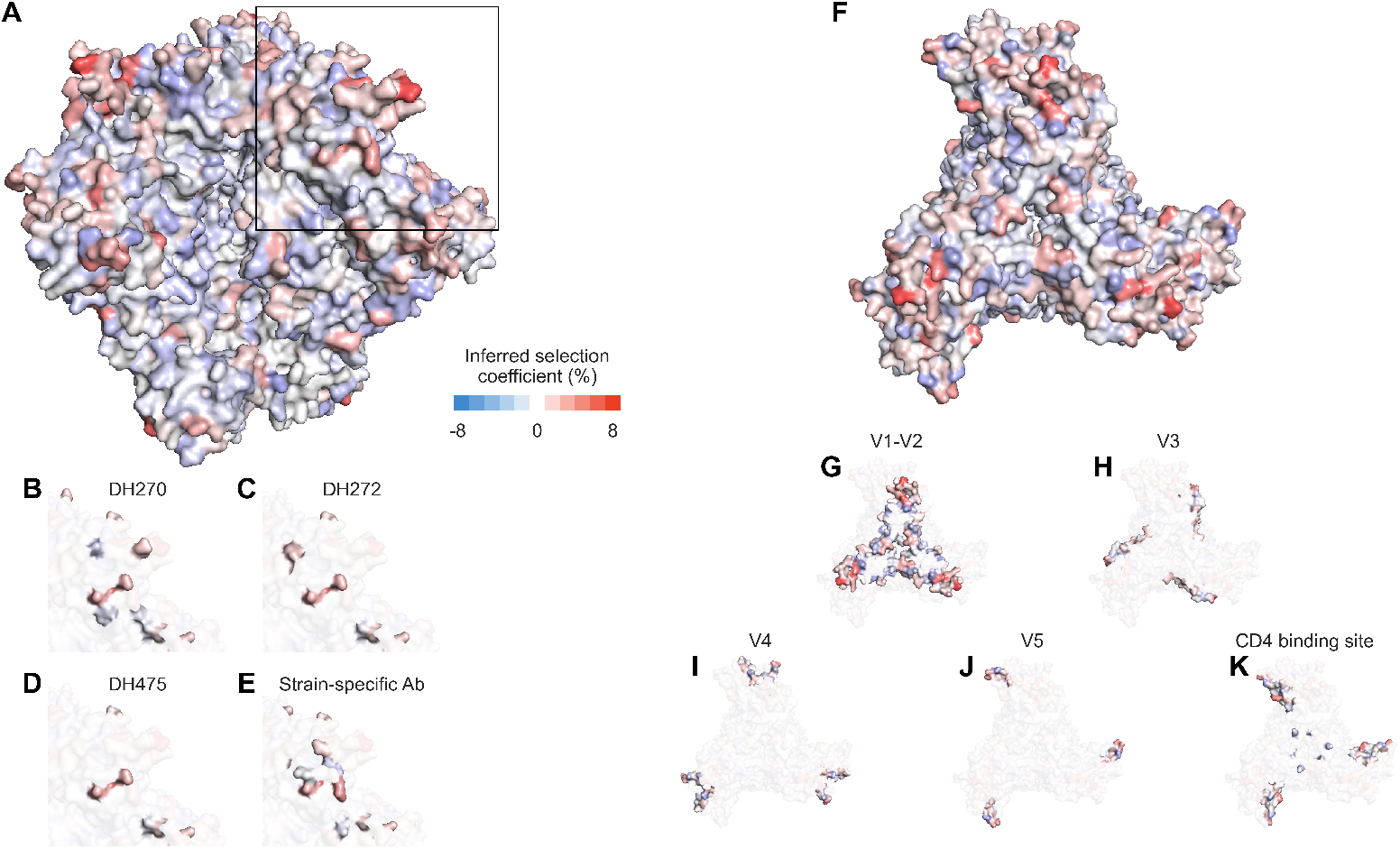
CH848 exhibits similar spatial patterns of selection coefficients; significantly strongly selected mutations are located at the apex, particularly on the edge of the Env protein. **A** and **B**, Side and top views of the Env protein with inferred selection coefficient values. The apex region is enriched with moderately and strongly beneficial mutations. **C-F**, Mutations that are presumably resistant to bnAbs and autologous nAbs are highlighted. The selection values in the autologous nAbs binding regions were slightly larger than those in the bnAbs binding regions. **G-K**, Top views highlight the individual variable regions.

**Supplementary Fig. 3.**
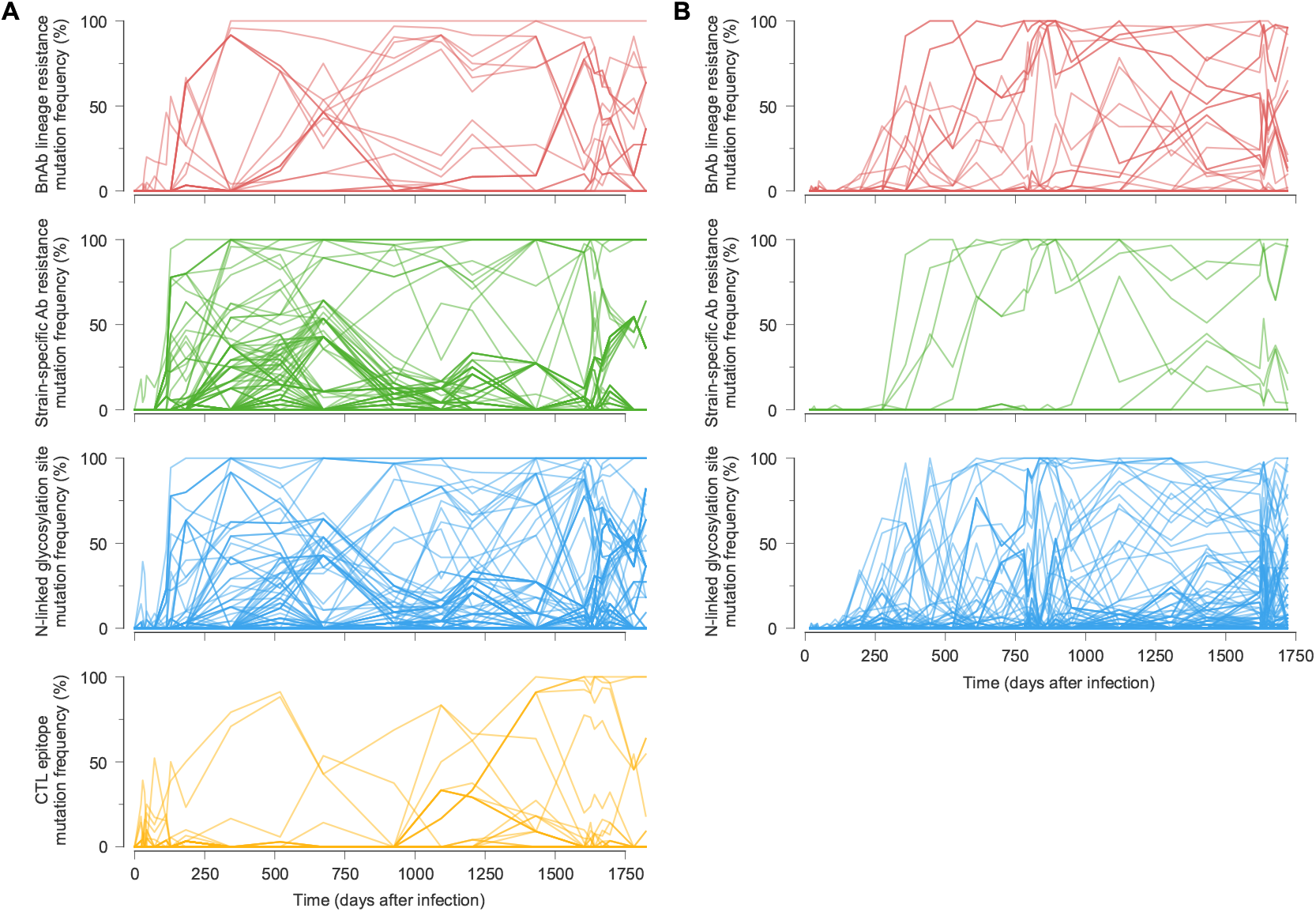
Frequencies of different types of HIV-1 mutations over time. CH505 (**A**) and CH848 (**B**) mutation frequencies over time. Mutation types are the same as in **Fig. 3** in the main text, but with all mutations affecting resistance to bnAb lineage antibodies grouped together.

**Supplementary Fig. 4.**
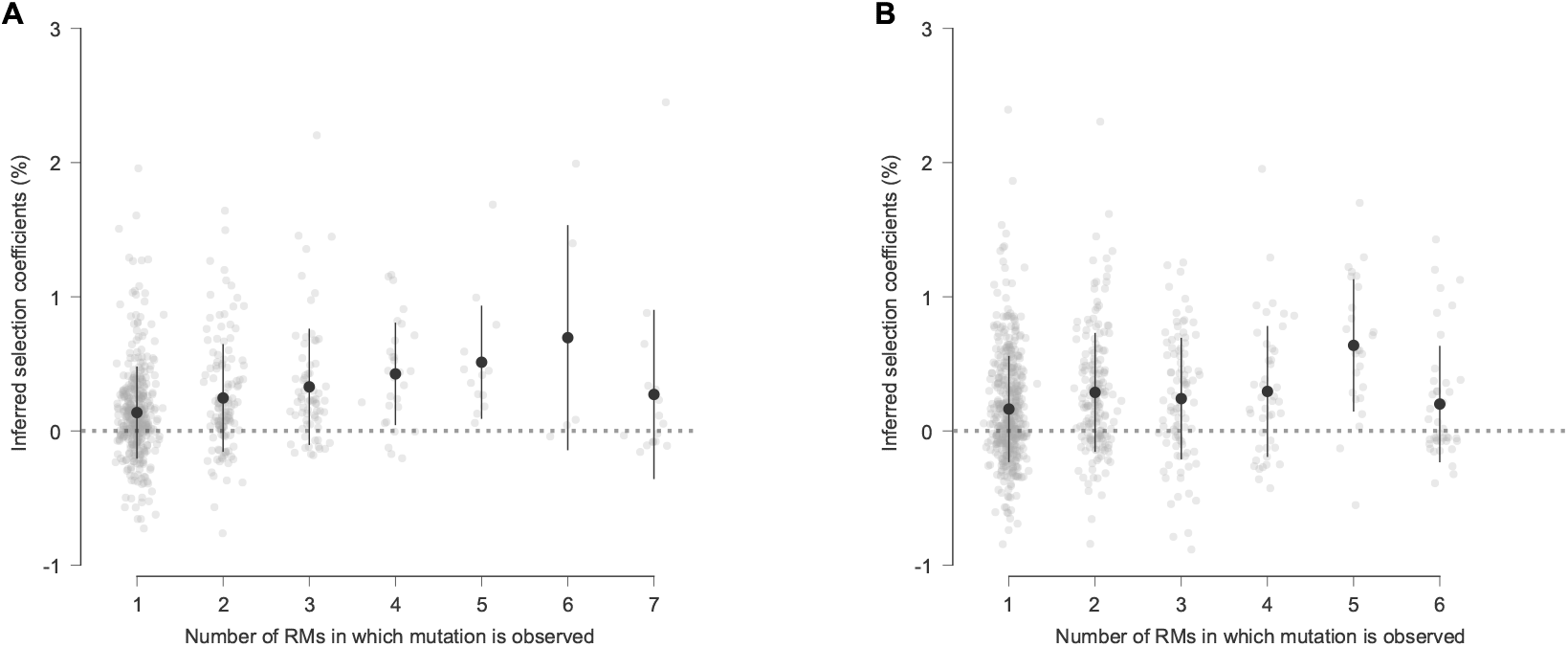
Weak association between the number of RMs in which a SHIV mutation was observed and its inferred fitness effect. Inferred selection coefficients for SHIV.CH505 (**A**) and SHIV.CH848 (**B**) mutations, sorted by the number of RMs in which the mutation was observed.

**Supplementary Fig. 5.**
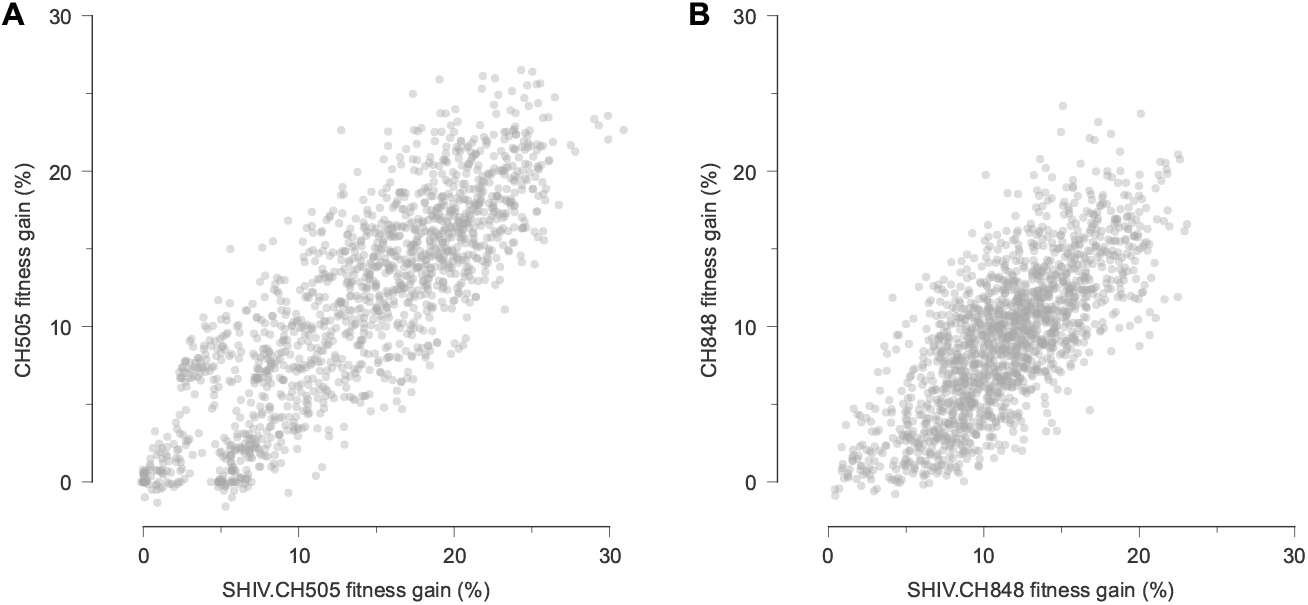
Broad similarity between HIV-1 and SHIV fitness landscapes with the same TF Env sequence. **A**, Fitness estimates for a sample of artificial Env sequences, obtained by independently shuffling observed amino acids in SHIV.CH505 sequences at each residue. This random sequence ensemble conserves single-residue frequencies, but not correlations between mutations. Even on these artificial sequences, the fitness estimates using a model trained on HIV-1 data strongly agree with the SHIV model (Pearson’s *r* = 0.84, *p <* 10^−20^). This implies that the similarity of the fitness landscapes is not confined to the specific genotypes observed in HIV-1 or SHIV evolution, but also extends to more distant sequences. **B**, Similar results also hold for fitness landscapes based on CH848 and SHIV.CH848 data (Pearson’s *R* = 0.74, *p <* 10^−20^).

**Supplementary Fig. 6.**
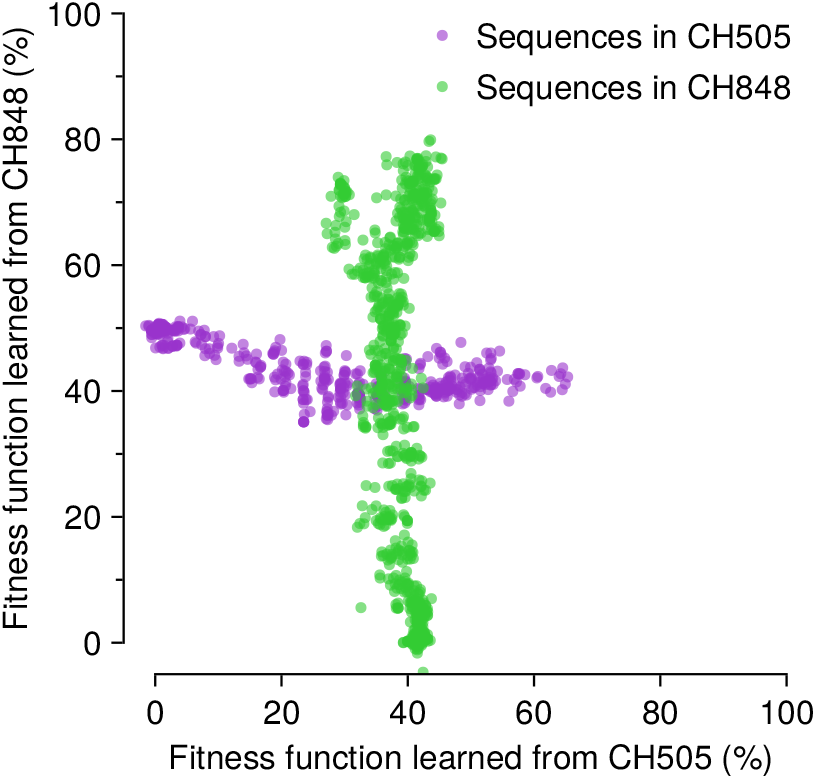
Little correlation in fitness values estimated from evolutionarily distant sequences. Comparison between estimated fitness values for HIV-1 sequences using fitness landscapes learned from CH505 and CH848 data. There is little correlation between the inferred landscapes. Furthermore, the CH505 landscape captures little variation in fitness for CH848 sequences (and vice versa). 199 mutations are shared between CH505 and CH848, among 868 and 1,406 total Env mutations, respectively.

**Supplementary Fig. 7.**
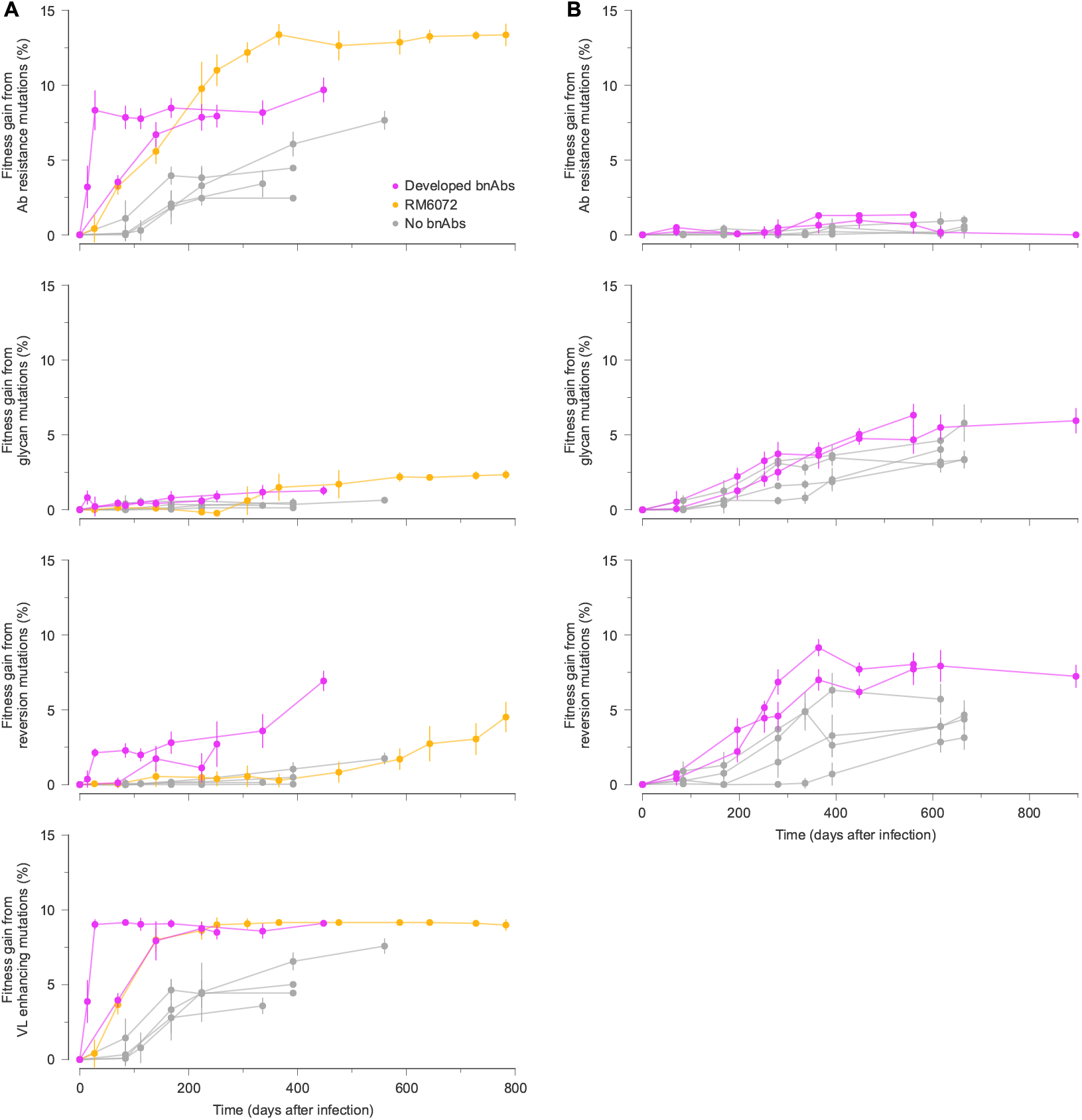
Contributions of different types of SHIV mutations to viral fitness gains over time. SHIV.CH505 (**A**) and SHIV.CH848 (**B**) mutations were grouped into four categories to assess their contributions to SHIV fitness gains over time. For SHIV.CH505, the mutations N334S, H417R, K302N, Y330H, N279D, and N130D are classified as viral load (VL) enhancing mutations, following ref. ^109^. For both SHIV.CH505 and SHIV.CH848, we then separated out the contributions of known antibody resistance mutations, including mutations that affect N-linked glycosylation motifs. We then computed the collective fitness contributions from subsets of mutations that affect N-linked glycosylation motifs that were not known to affect resistance to specific antibodies, and reversions to the HIV-1 subtype consensus sequence. Each mutation appears in only one category in this figure, sorted in the order above. For example, a mutation that affects an N-linked glycosylation motif and which is a reversion to the subtype consensus sequence, but which has not been established to affect resistance to a specific antibody, would have its contribution to fitness counted in the glycan category.

**Supplementary Fig. 8.**
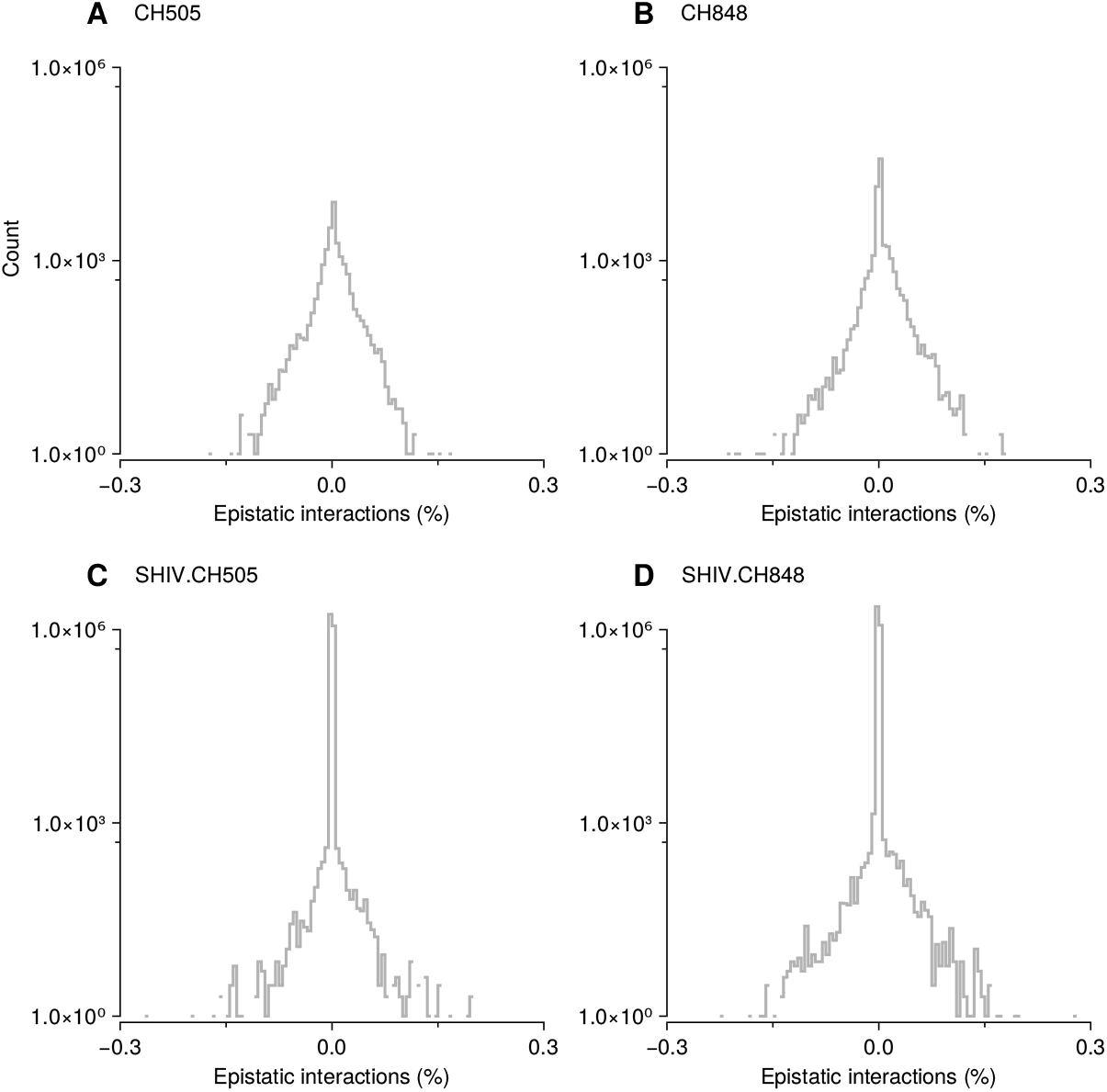
Most of the inferred epistatic interactions concentrate around zero, with only a small fraction exhibiting moderately larger absolute values. represents the distribution of inferred epistatic interactions forCH505 (**A**), CH848 (**B**), SHIV.CH505 (**C**), and SHIV.CH848 (**D**).

**Supplementary Fig. 9.**
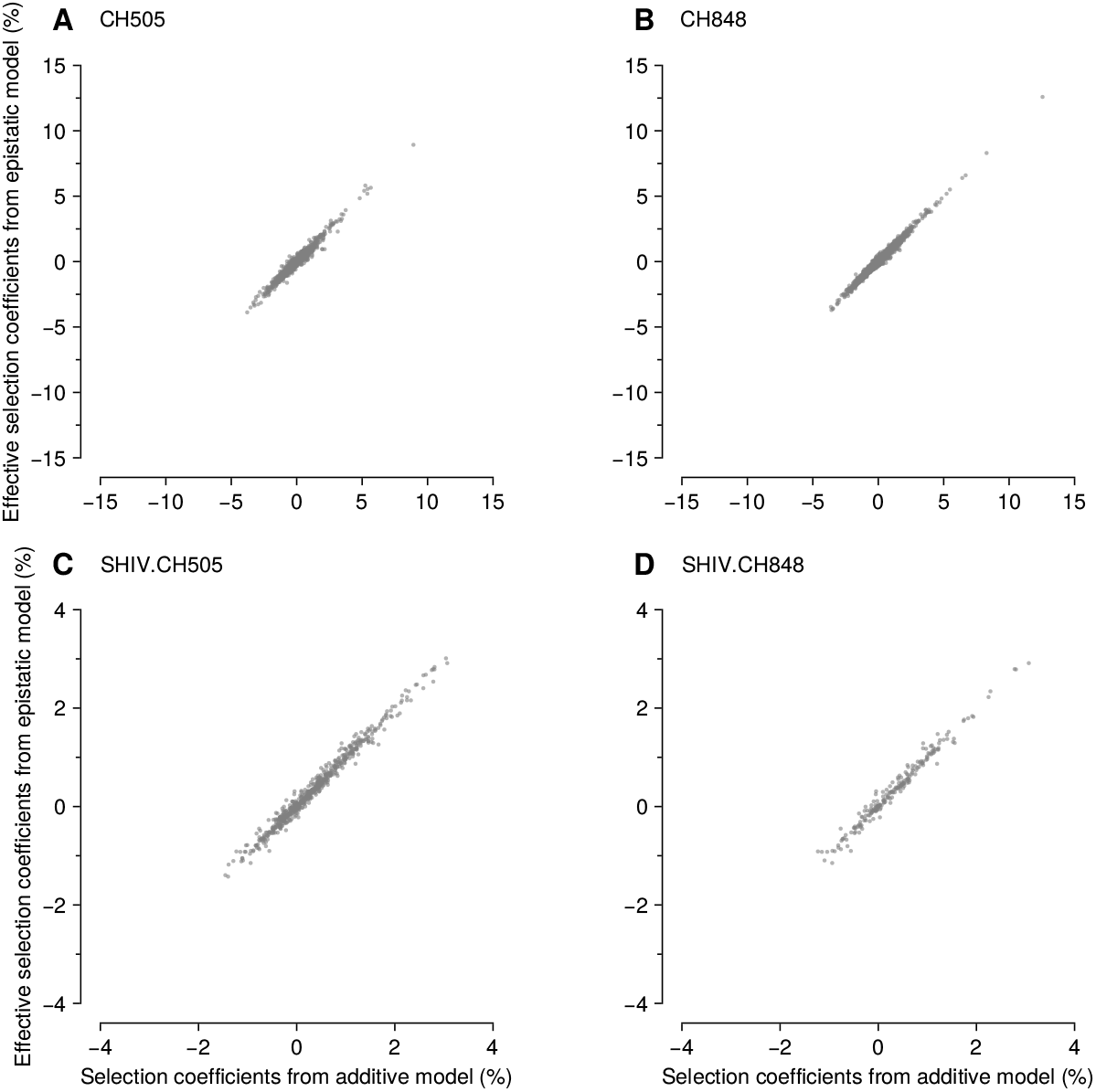
Effective selection coefficients obtained from the epistatic model align well with the selection coefficients in the additive model. We compared selection coefficients from the additive model analyzed throughout the manuscript to the effective selection coefficients from the epistatic model for CH505 (**A**) and CH848 (**B**). Intuitively, the effective selection coefficient is defined as the average difference in fitness when a particular mutation is replaced by the TF nucleotide/amino acid. For definiteness, let *g* represent an arbitrary sequence and let *g*^*i\**^ represent a sequence that is identical to *g*, except that the nucleotide/amino acid at site *i* has been replaced by the TF one. The effective selection coefficient is then defined as 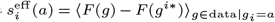, where the average runs over all the sequences in the data set that have mutant allele *a* at site *i*. Similarly, we compared selection from the additive and epistatic models for SHIV.CH505 (**C**) and SHIV.CH848 (**D**).

**Supplementary Fig. 10.**
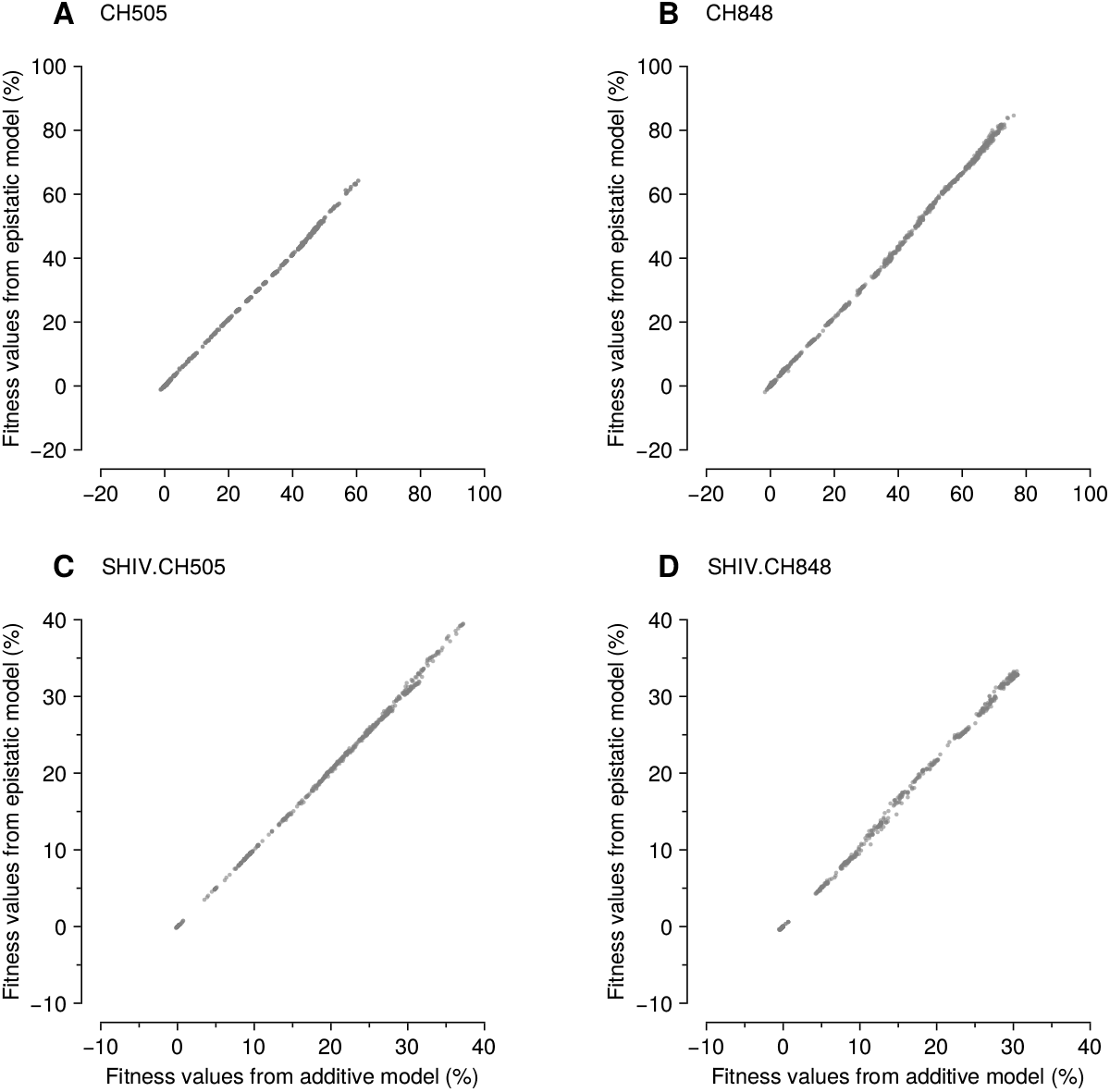
Fitness values are consistent between the additive and epistatic models. Compare the fitness values obtained from the additive and epistatic models for CH505 (**A**), CH848 (**B**), SHIV.CH505 (**C**), and SHIV.CH848 (**D**), respectively.

**Supplementary Fig. 11.**
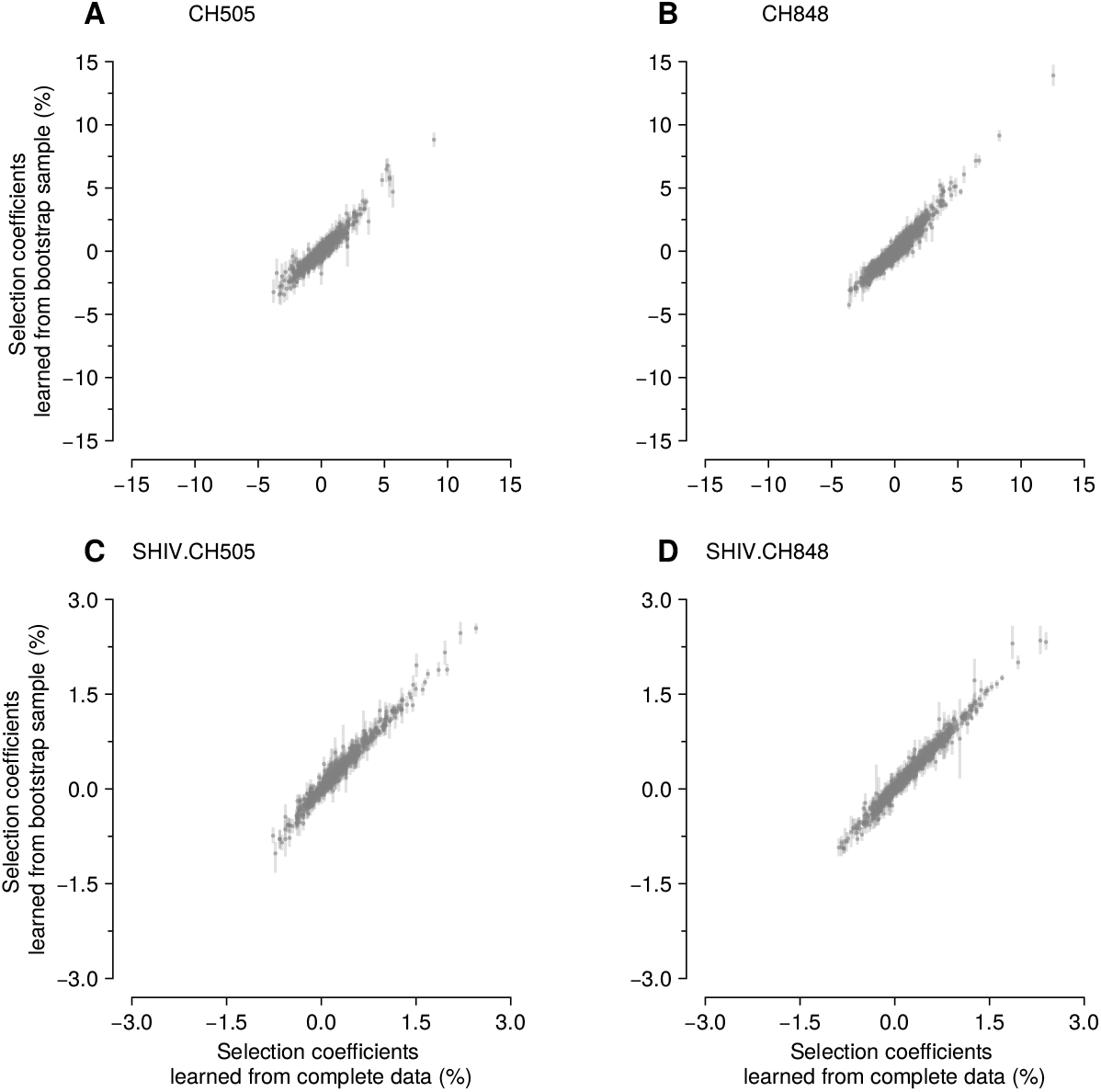
Selection coefficients are robust to finite sampling noise. Selection coefficients from the full and bootstrap-resampled data are compared for CH505 (**A**), CH848 (**B**), SHIV.CH505 (**C**), and SHIV.CH848 (**D**). Each point and error bar represents the mean and confidence interval, respectively, based on 10 independently inferred selection coefficients from bootstrap samples. The bootstrap samples are obtained by uniformly resampling the same number of sequences from the sequence ensemble for each subject at each time point.

**Supplementary Table 1.**
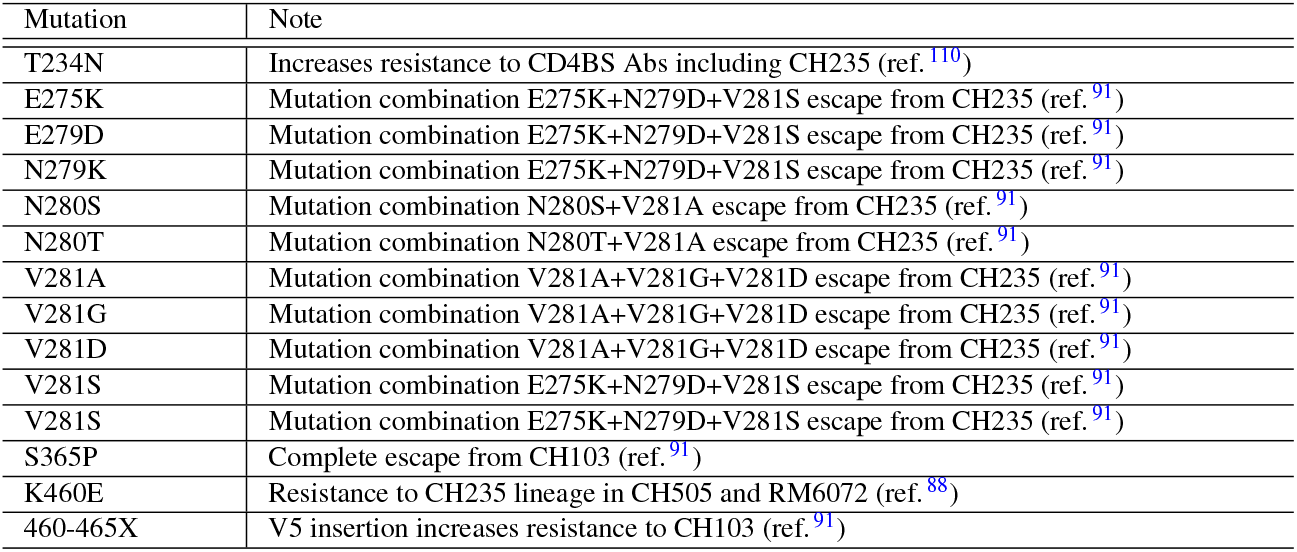
CH103 and CH235 resistance mutations. List of mutations considered to be CH103 or CH235 resistance mutations, and supporting references. Here “X” refers to any amino acid.

**Supplementary Table 2.**
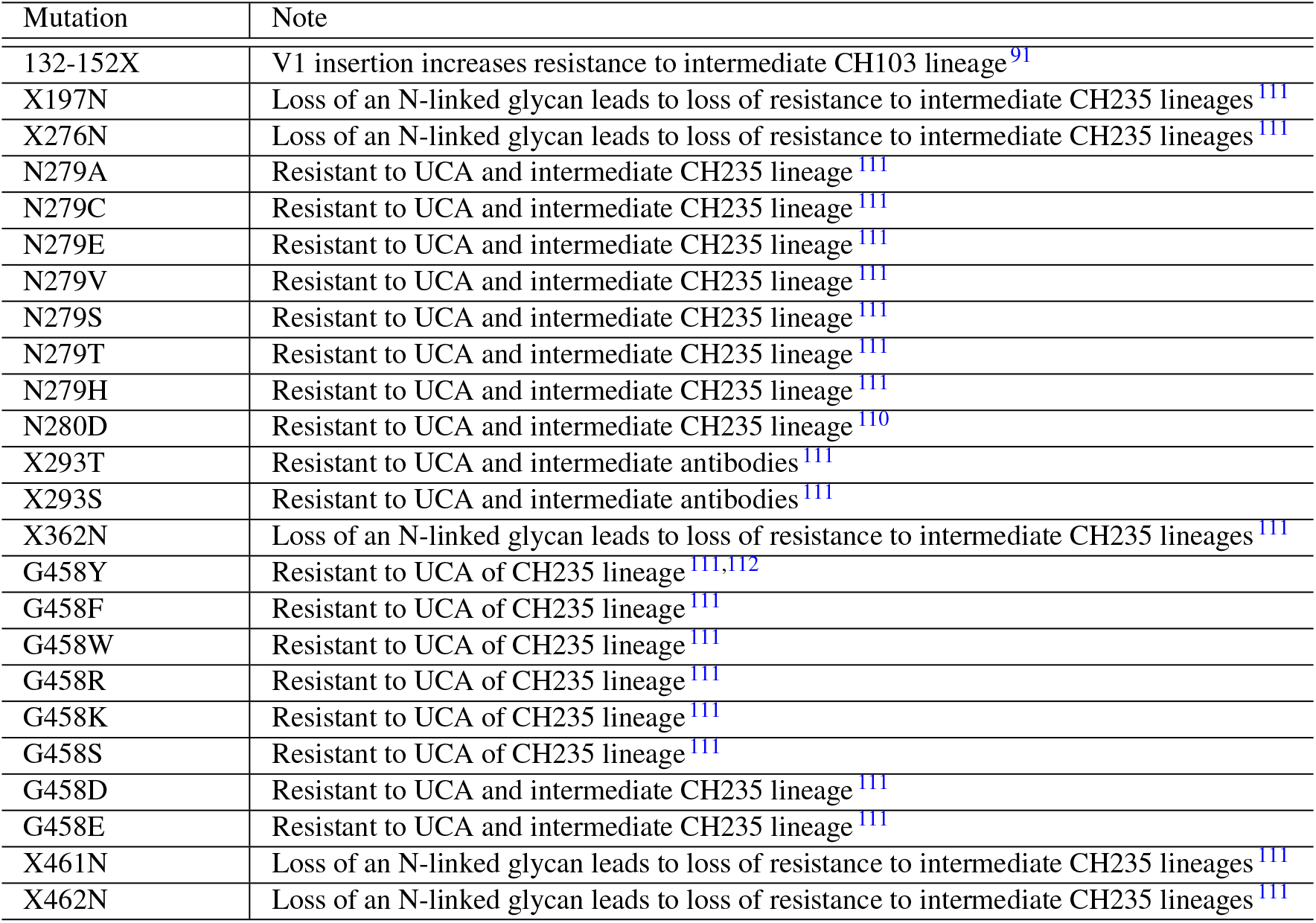
Strain-specific antibody resistance mutations in CH505. List of mutations considered to be strain-specific antibody resistance mutations and supporting references. Here “X” refers to any amino acid.

**Supplementary Table 3.**
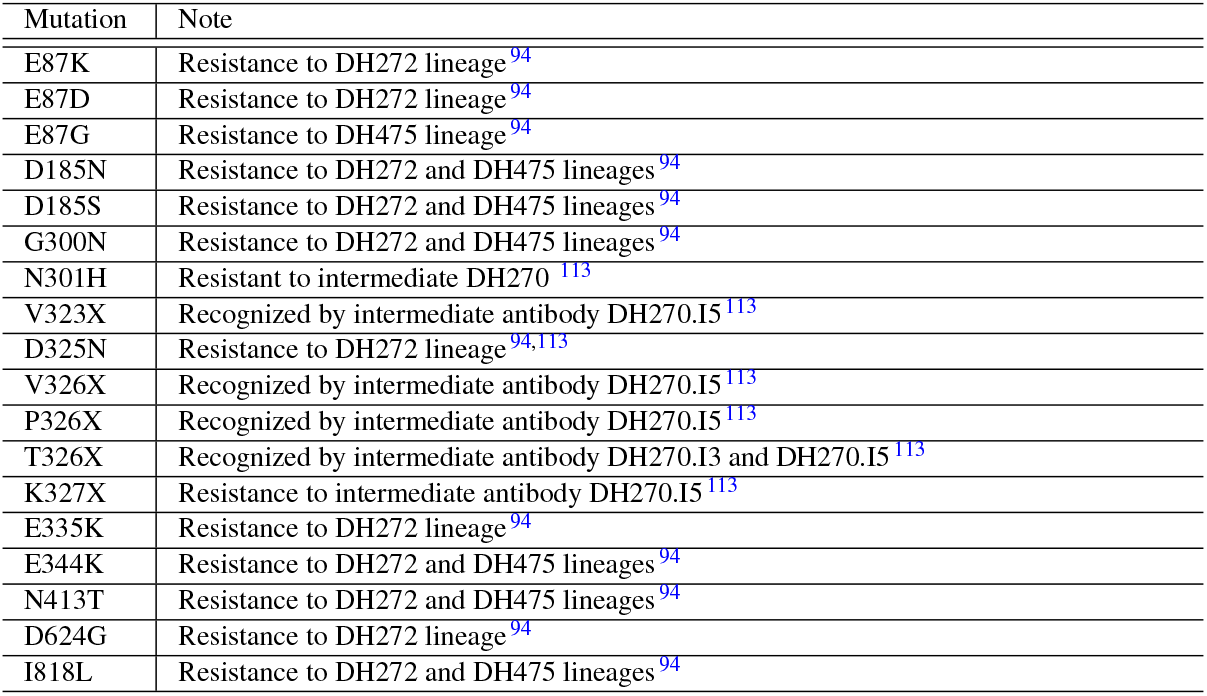
DH272, DH475, and strain-specific antibody resistance mutations in CH848. List of mutations considered to be DH272, DH475, and strain-specific antibody resistance mutations, and supporting references. Here “X” refers to any amino acid.

| Mutation | Note |
| --- | --- |
| E87K | Resistance to DH272 lineage<sup>94</sup> |
| E87D | Resistance to DH272 lineage<sup>94</sup> |
| E87G | Resistance to DH475 lineage<sup>94</sup> |
| D185N | Resistance to DH272 and DH475 lineages<sup>94</sup> |
| D185S | Resistance to DH272 and DH475 lineages<sup>94</sup> |
| G300N | Resistance to DH272 and DH475 lineages<sup>94</sup> |
| N301H | Resistant to intermediate DH270<sup>113</sup> |
| V323X | Recognized by intermediate antibody DH270.I5<sup>113</sup> |
| D325N | Resistance to DH272 lineage<sup>94,113</sup> |
| V326X | Recognized by intermediate antibody DH270.I5<sup>113</sup> |
| P326X | Recognized by intermediate antibody DH270.I5<sup>113</sup> |
| T326X | Recognized by intermediate antibody DH270.I3 and DH270.I5<sup>113</sup> |
| K327X | Resistance to intermediate antibody DH270.I5<sup>113</sup> |
| E335K | Resistance to DH272 lineage<sup>94</sup> |
| E344K | Resistance to DH272 and DH475 lineages<sup>94</sup> |
| N413T | Resistance to DH272 and DH475 lineages<sup>94</sup> |
| D624G | Resistance to DH272 lineage<sup>94</sup> |
| I818L | Resistance to DH272 and DH475 lineages<sup>94</sup> |

**Supplementary Table 4.** Biological effects of the HIV-1 mutations inferred to be the most beneficial in CH505. The table lists the 25 mutations in CH505 with the highest (i.e., most beneficial) selection coefficients, among over 2500 mutations in total. All of the top mutations are nonsynonymous. Mutations associated with immune evasion, changes in glycosylation, and reversions to the subtype C consensus sequence are frequently observed among these top mutations.

| Rank | Mutation (DNA) | Mutation (AA) | Date | Region | Selection coefficient (%) | Note |
| --- | --- | --- | --- | --- | --- | --- |
| 1 | A7225G | N334S | 113 | Near V3 Loop | 8.92 | Reversion, N-linked glycan shift, resistance to DH950 antibodies<sup>114</sup> |
| 2 | T7066A | V281D | 183 | CD4bs/Loop D | 5.66 | Resistant to intermediate CH235 lineage antibodies<sup>91</sup>, combination with N280S or N280T gives resistance to mature CH235 (ref.<sup>91</sup>) |
| 3 | G7776A | V518M | 1093 | Fusion Peptide | 5.42 |  |
| 4 | C6709T | T162I | 129 | V2 Loop | 5.40 | N-linked glycan hole |
| 5 | G6682A | G153E | 71 | V1 Loop | 5.24 | Reversion |
| 6 | C7462A | T413N | 129 | V4 Loop | 5.16 | N-linked glycan shift |
| 7 | C7468A | T415K | 71 | V4 Loop | 4.82 | Putative CD8<sup>+</sup> T cell escape mutation<sup>91</sup> |
| 8 | A7130C | K302N | 183 | V3 Loop | 3.74 | Reduces neutralization of DH950-like Abs<sup>114</sup> |
| 9 | G7769A | M515I | 1603 | Fusion Peptide | 3.57 | Reversion |
| 10 | A7614G | N464D | 183 | V5 Loop | 3.46 | Reversion |
| 11 | C7462T | T413I | 23 | V4 Loop | 3.42 | Putative CD8<sup>+</sup> T cell escape mutation<sup>91</sup> |
| 12 | G7776T | V518L | 1640 | Fusion Peptide | 3.37 |  |
| 13 | T7212C | Y330H | 129 | V3 Loop | 3.28 | Reversion, reduces neutralization of DH950-like Abs<sup>114</sup> |
| 14 | G6661A | S146N | 129 | V1 Loop | 3.14 | Reversion, N-linked glycan shield |
| 15 | G7453A | S410N | 23 | V4 Loop | 3.08 | Putative CD8<sup>+</sup> T cell escape mutation<sup>93</sup> |
| 16 | G6634T | S137I | 183 | V1 Loop | 3.00 | N-linked glycan hole |
| 17 | G6634C | S137A | 129 | V1 Loop | 2.82 | Reversion, N-linked glycan hole |
| 18 | T6588A | L122M | 183 | Near CD4bs | 2.81 |  |
| 19 | C7456A | T411K | 29 | V4 Loop | 2.75 | Reversion, N-linked glycan hole, and putative CD8<sup>+</sup> T cell escape mutation<sup>93</sup> |
| 20 | A6596C | P124P | 183 | CD4bs | 2.73 |  |
| 21 | A7474G | H417R | 23 | V4 Loop | 2.63 | Putative CD8<sup>+</sup> T cell escape mutation<sup>93</sup> |
| 22 | A7601dG | N459D | 1603 | CD4bs | 2.61 | N-linked glycan shield, located in the CH235 antibody binding region<sup>93</sup> |
| 23 | A7056T | T278S | 1603 | Loop D | 2.59 | Located in the CH235 antibody binding region<sup>93</sup> |
| 24 | G6630A | A136T | 113 | V1 Loop | 2.59 | Reversion |
| 25 | A8143G | N640S | 1780 | Near Glycosite | 2.55 |  |

**Supplementary Table 5.** Biological effects of the HIV-1 mutations inferred to be the most beneficial in CH848. This table is analogous to **Supplementary Table 4**, but for CH848. Unlike for CH505, longer sequences were available for CH848, including genes beyond Env. We observed multiple beneficial mutations in Nef and Env.

| Rank | Mutation (DNA) | Mutation (AA) | Date | Region | Selection co-efficient (%) | Note |
| --- | --- | --- | --- | --- | --- | --- |
| 1 | T9043G | S83A | 49 | Nef | 12.55 | Reversion |
| 2 | A8988cT | E65D | 38 | Nef | 8.28 |  |
| 3 | G8911A | E39K | 38 | Nef | 6.68 | Reversion |
| 4 | A8988bG | E65G | 38 | Nef | 6.42 |  |
| 5 | A8988cC | E65D | 43 | Nef | 5.48 |  |
| 6 | G6640A | S139N | 358 | V1 Loop | 5.22 | Reversion, N-linked glycan shift, presence of N-linked glycan at site 138 decreases neutralization by the earlier DH270 UCA and removes glycan at site 137 (ref. <sup>114</sup>) |
| 7 | G6679A | G152E | 38 | V1 Loop | 4.83 |  |
| 8 | C9044G | A84G | 78 | Nef | 4.70 | Reversion |
| 9 | A8915G | H41R | 38 | Nef | 4.48 | Reversion |
| 10 | A8083T | K620M | 808 | gp41 | 4.44 |  |
| 11 | A8988cG | E65E | 38 | Nef | 4.35 |  |
| 12 | G7123A | S300N | 700 | V3 Loop | 4.06 | Reversion, confers resistance to DH272 and DH475 (ref. <sup>94</sup>) |
| 13 | G6640T | S139V | 445 | V1 Loop | 3.92 | N-linked glycan hole |
| 14 | A7310C | K362N | 780 | V1 Loop | 3.87 | N-linked glycan shield |
| 15 | A8083C | K620T | 780 | gp41 | 3.84 |  |
| 16 | A8912C | E40A | 32 | Nef | 3.83 |  |
| 17 | A8895G | A34A | 49 | Nef | 3.77 | Reversion |
| 18 | A8912T | E40V | 43 | Nef | 3.74 |  |
| 19 | G6640C | S139T | 1432 | V1 Loop | 3.73 | N-linked glycan shield |
| 20 | A6621C | N133H | 1120 | V1 Loop | 3.66 | N-linked glycan hole |
| 21 | G8911C | E39Q | 49 | Nef | 3.64 |  |
| 22 | C7066T | A281V | 1651 | CD4BS | 3.62 | Reversion |
| 23 | A6912G | N230D | 194 | Glycosite | 3.57 | N-linked glycan hole, amino acid N is sensitive to DH270.5 (ref. <sup>110</sup>) |
| 24 | Δ8911G | Δ39E | 78 | Nef | 3.57 |  |
| 25 | A7614G | T464E | 445 | V5 Loop | 3.39 | Reversion, N-linked glycan hole |

**Supplementary Table 6.**
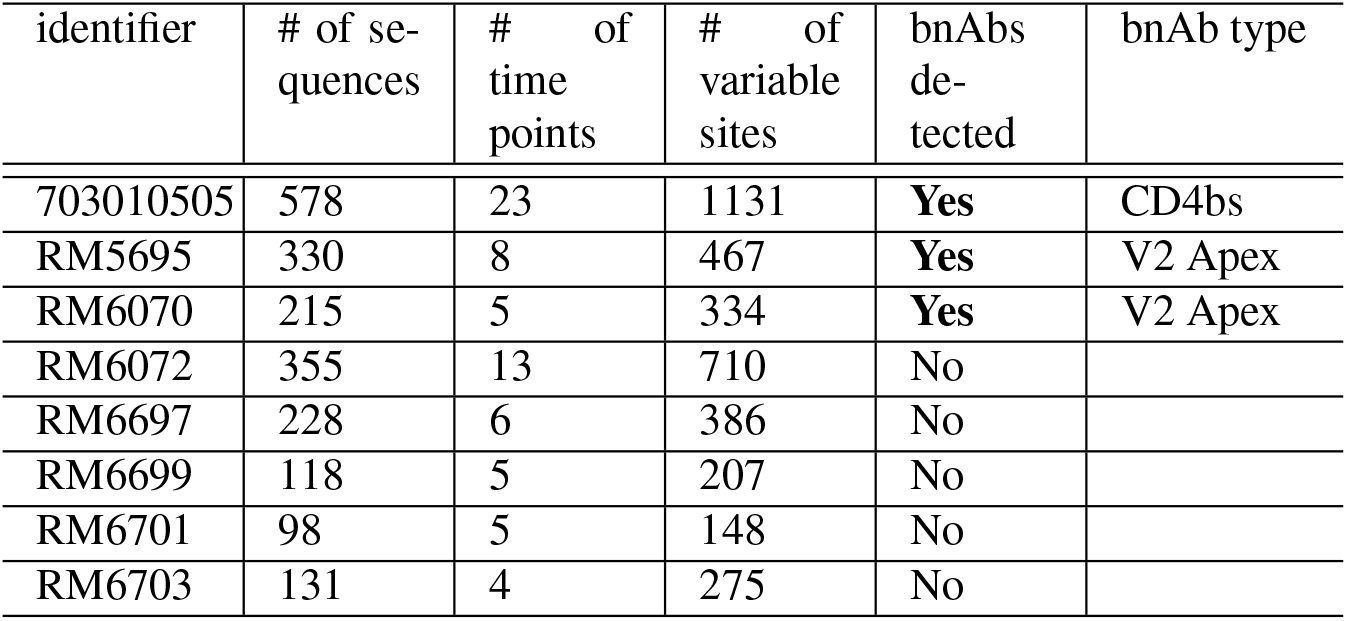
CH505 and SHIV.CH505 sequence statistics. The number of sequences and time points listed here are given after filtering. Variable sites are defined as sites where at least two different nucleotides (including deletions) were observed.

| identifier | # of sequences | # of time points | # of variable sites | bnAbs detected | bnAb type |
| --- | --- | --- | --- | --- | --- |
| 703010505 | 578 | 23 | 1131 | **Yes** | CD4bs |
| RM5695 | 330 | 8 | 467 | **Yes** | V2 Apex |
| RM6070 | 215 | 5 | 334 | **Yes** | V2 Apex |
| RM6072 | 355 | 13 | 710 | No |  |
| RM6697 | 228 | 6 | 386 | No |  |
| RM6699 | 118 | 5 | 207 | No |  |
| RM6701 | 98 | 5 | 148 | No |  |
| RM6703 | 131 | 4 | 275 | No |  |

**Supplementary Table 7.** CH848 and SHIV.CH848 sequence statistics. The format of this table is the same as in **Supplementary Table 6**.

| identifier | # of sequences | # of time points | # of variable sites | bnAbs detected | bnAb type |
| --- | --- | --- | --- | --- | --- |
| 703010848 | 1205 | 31 | 2694 | **Yes** | V3-glycan |
| RM6163 | 261 | 8 | 467 | **Yes** | V3-glycan |
| RM6167 | 349 | 10 | 657 | **Yes** | V3-glycan |
| RM6700 | 260 | 7 | 568 | No |  |
| RM6713 | 281 | 7 | 567 | No |  |
| RM6714 | 284 | 8 | 516 | No |  |
| RM6720 | 193 | 8 | 340 | No |  |

**Supplementary Table 8.** Biological effects of strongly selected SHIV.CH505 mutations. Five out of the top six mutations have been shown to increase viral load in RMs ^109^. Selection coefficients and ranks are based on the joint evolutionary model across SHIV.CH505-infected RMs. These analyses focus on common mutations observed in three or more RMs.

| Rank | Mutation (AA) | Region | Selection coefficient (%) | Note |
| --- | --- | --- | --- | --- |
| 1 | N334S | Near V3 Loop | 2.44 | N-linked glycan shift, enhanced viral replication<sup>109</sup>, reduced early plasma neutralization<sup>88</sup> |
| 2 | Y330H | V3 Loop | 1.99 | Improved virus entry and replication<sup>109</sup> |
| 3 | N279D | CD4BS | 1.69 | Enhanced replication<sup>109</sup>, reduced early plasma neutralization, conferred resistance to strain-specific CD4bs nAbs DH650 in RM6070 (ref.<sup>88</sup>) |
| 4 | K462E | V5 Loop | 1.40 |  |
| 5 | H356N | Glycosite | 1.16 |  |
| 6 | K302N | V3 Loop | 1.15 | Enhanced replication and affects virus entry, conferred resistance to strain-specific V3 nAbs, DH647 and DH648 (ref. <sup>109</sup>) |
| 7 | G153E | V1 Loop | 1.12 |  |
| 8 | N130D | Near V1 Loop | 0.99 | Reduced early plasma neutralization<sup>88</sup> |
| 9 | T234N | Glycosite | 0.91 | N-linked glycan shield, reduced early plasma neutralization, conferred resistance to strain-specific CD4bs nAbs DH650 in RM6070 (ref. <sup>88</sup>) |
| 10 | H417R | V4 Loop | 0.87 | Enhanced replication<sup>109</sup>, putative CD8<sup>+</sup> T cell escape mutation<sup>88,109</sup> |

**Supplementary Table 9.** Biological effects of strongly selected SHIV.CH848 mutations. Selection coefficients and ranks are based on the joint evolutionary model across SHIV.CH848-infected RMs.

| Rank | Mutation (AA) | Region | Selection coefficient (%) | Note |
| --- | --- | --- | --- | --- |
| 1 | P179L | V2 Loop | 1.95 |  |
| 2 | N413T | V4 Loop | 1.70 | N-linked glycan hole, loss of N413 alters sensitivity to early nAbs in RM6167 (ref. <sup>88</sup>) |
| 3 | N413S | V4 Loop | 1.43 | N-linked glycan hole, aids in antibody escape <sup>88</sup> |
| 4 | D137S | V1 Loop | 1.29 |  |
| 5 | N462T | V4 Loop | 1.29 | N-linked glycan shift |
| 6 | S407N | V4 Loop | 1.22 |  |
| 7 | R336E | Near V3 Loop | 1.20 | Mutations at site 336 observed in all RMs, R336G reduced sensitivity to plasma in RM6167 (ref. <sup>88</sup>) |
| 8 | P179S | V2 Loop | 1.18 |  |
| 9 | V142fA | V1 Loop | 1.15 |  |
| 10 | N462D | V4 Loop | 1.15 | N-linked glycan hole |

**Supplementary Table 10.** Selective advantage of mutations that confer resistance to antibodies in SHIV.CH505. Identification of resistance mutations was performed by Roark and colleagues ^88^.

| Mutation | RM5695<br>(%) | RM6072<br>(%) | RM6701<br>(%) | RM6699<br>(%) | RM6697<br>(%) | RM6070<br>(%) | RM6703<br>(%) | Note |
| --- | --- | --- | --- | --- | --- | --- | --- | --- |
| N130D | 1.5 | 1.1 |  |  | 1.0 | 0.90 | 0.87 | Confers resistance to V3 mAbs DH647 and DH648 in RM6072 |
| R166K | 0.49 |  |  |  |  | 1.77 |  | Confers resistance to V2 apex bnAbs RHA1 in RM5695 |
| R166S | 0.7 |  |  |  |  |  |  | Confers resistance to V2 apex bnAbs RHA1 in RM5695 |
| R169K | 0.43 | -0.1 |  |  |  | -0.04 |  | Confers resistance to V2 apex bnAbs RHA1 in RM5695 |
| T234N | 0.36 | 4.5 |  |  | 0.34 | 1.1 |  | Confers resistance to CD4bs mAbs DH650 in RM6072 |
| N279K | 1.2 | 0.013 | -0.32 | 0.44 | -0.19 | -0.3 |  | Confers resistance to CD4bs mAbs DH650 in RM6072 |
| V281I | 0.14 | -0.03 |  | 0.59 |  | 1.4 |  | Confers resistance to V3 mAbs DH647 and DH648 in RM6072 |
| K302N | 6.19 | 0.59 |  |  | 0.76 |  |  | Confers resistance to V3 mAbs DH647 and DH648 in RM6072 |
| Y330H | 6.5 | 3.0 | 0.03 | 1.1 | 0.11 | -0.56 | 1.4 | Confers resistance to V3 mAbs DH647 and DH648 in RM6072 |
| N334S | 7.2 | 4.6 | 1.8 | 1.6 | 1.4 | 3.1 | 2.4 | Confers resistance to V3 mAbs DH647 and DH648 in RM6072 |

**Supplementary Table 11.** Selective advantage of mutations that confer resistance to antibodies in SHIV.CH848. Identification of resistance mutations was performed by Roark and colleagues ^88^.

| Mutation | **RM6163**<br>(%) | **RM6167**<br>(%) | **RM6700**<br>(%) | **RM713**<br>(%) | **RM6714**<br>(%) | **RM6720**<br>(%) | **Note** |
| --- | --- | --- | --- | --- | --- | --- | --- |
| N332S |  | -0.19 |  |  |  |  | Mutations at or near N332 to 334 can modulate sensitivity to neutralization |
| R336G | 0.62 | 0.4 | 0.94 | 0.24 | 0.12 | 0.98 | Led to a reduction in neutralization by autologous nAb in RM6167 |

**Supplementary Table 12.** List of selected sites using LASSIE in SHIV.CH505. This analysis was performed by Roark et al. ^88^.

| HXB2 Site |
| --- |
| N130 |
| T234 |
| N279 |
| V281 |
| N300 |
| K302 |
| Y330 |
| N334 |
| K347 |
| H355 |
| T413 |
| H417 |
| G471 |
| G620 |
| E649 |

**Supplementary Table 13.** List of selected sites using LASSIE in SHIV.CH848. This analysis was performed by Roark et al. ^88^.

| **HXB2 Site** |
| --- |
| N130 |
| N289 |
| S291 |
| G300 |
| R336 |
| Q337 |
| K340 |
| E344 |
| R363 |
| A372 |
| T411 |
| N413 |
| T467 |
| H651 |
| K658 |
| R683 |

## Notes

https://github.com/bartonlab/paper-HIV-coevolution

### Competing Interest Statement

The authors have declared no competing interest.

### Summary of Updates

We analyzed the robustness of our results to variation in the fitness model (i.e., epistatic versus purely additive) and noise due to finite sampling. We have also revised the text to improve clarity and more fully discuss both the limitations and potential implications of this work.

